# Measuring Perfusion Pressure and Flow Resistance in a Microfluidic Device Using an External Optical System

**DOI:** 10.1101/2025.11.03.686291

**Authors:** Matthew C. Coughlin, Marie A. Floryan, Giovanni S. Offeddu, Mark F. Coughlin

**Affiliations:** Department of Mechanical and Industrial Engineering, Northeastern University, Boston, MA 02115; Department of Mechanical Engineering, Massachusetts Institute of Technology, Cambridge, MA 02139; Department of Biological Engineering, Massachusetts Institute of Technology, Cambridge, MA 02139

## Abstract

The pathology of human diseases are now investigated using microphysiological systems (MPS) supporting vascular structures. Efforts to increase physiological relevance of these platforms have centered on the incorporation of organ-specific cellular and non-cellular constituents. However, tissue-specific cellular constituents must experience appropriate physical forces to faithfully replicate physiological function. Quantification of physical forces in MPS has received little attention. The goal of this study was to establish a simple and robust system capable of interfacing with existing pumps to quantitatively characterize the flow delivered to a MPS. The system assessed both the fluid pressure driving flow through a microphysiological platform and the resistance to flow of glass capillary tubes or a model vascular network. The system showed excellent qualitative and quantitative agreement with resistance values measured by a hydrostatic approach and predicted for laminar flow through a smooth capillary tube. Importantly, the system is optically-based without sensors contacting the circulating fluid making it ideally-suited for long-term biological studies where sterility is paramount. Benchmarking experiments were supplemented with measurements of driving pressure and flow resistance from vascular structures within a MPS in a humidified incubator. Vascular resistance measurements were consistent with published results obtained from similar microvascular networks.

## Introduction

An important experimental tool that bridges the gap between cell culture and animal models is the engineered microphysiological system (MPS). A typical MPS consists of a combination of cellular and noncellular biological materials within an engineered environment. A MPS may be as simple as a pair of media channels flanking a single channel containing a hydrated gel infused with endothelial cells that will spontaneously self-assemble into three-dimensional vasculature networks [1-3]. Vascularized MPS showed immediate promise as a novel tool to study circulatory pathologies [cf. 4]. More sophisticated MPS exhibiting organ-specific functionality emerged nearly exclusively through the modulation of the cellular constituents [5]. However, a powerful determinant of organ-specific functionality derives from physical cues originating in the tissue environment [6, 7]. Eliciting enhanced physiological relevance and tissue-specific physiology requires addressing the difficult and largely overlooked engineering problem of controlling the physical environment within a MPS.

The function of vascular cells is profoundly influenced by intravascular and interstitial flow [8]. A pump was developed to recirculate a small media volume at physiologically relevant pressures and flows in MPS [9]. The pump consisted of a pumping chamber, a pair of check valves, and a pair of fluid capacitors. The main function of the pump was to maintain a pressure difference between the capacitors. Using the pump to regulate fluid forces in a MPS required feedback. In particular, the flow through a MPS is determined by both the driving pressure of the pump and flow resistance offered of the system. Thus, unlocking the potential to influence physiological function in MPS requires quantitative characterization of the physical performance of the pump and flow resistance of the MPS.

An optically-based system was developed to characterize both the driving pressure of the pump and the flow resistance of the MPS. Performance was quantified by measuring the pressure difference generated at the pump to drive flow. Resistance was quantified by measuring pressure transients. Functionality of the optical system was first demonstrated using glass capillary tubes with known flow resistances. Flow resistance was also measured in model bifurcating network that better replicated flow through a branching microvasculature. Importantly, the system quantitatively characterized both pump performance and flow resistance without a sensor contacting the circulating fluid. This makes the system ideally suited for long-term perfusion experiments of MPS supporting vascular structures that must be maintained under sterile conditions. The capability to function in a sterile environment was demonstrated by monitoring pump pressure and flow resistance of a MPS supporting a microvascular network maintained in a humidified incubator over ∼48 h. Taken together, this system offers critical functionality allowing quantification of key parameters responsible for the physical cues originating from intravascular and interstitial fluid flow that ultimately induce physiological relevance and organ-specific functionality to MPS.

## Methods

### Pump Fabrication

Pump assembly followed a published protocol [9] with modifications for the optical system. Pumps were fabricated from PDMS and activator at a ratio of 10:1 (SYLGARD 184 Silicone Elastomer, Dow Silicones Corporation) poured into a mold fabricated in-house and cured overnight at 60°C (Fig. 1A). The mold included narrow ridges that produced grooves in the cured PDMS that accepted a razor blade to facilitate accurate sizing of upper and lower pump slabs (Fig. 1B and C). Key features incorporated into the mold include narrow channels, a pumping chamber, a pair of check valves, a measurement capacitor, a reference capacitor, an inlet port, and an outlet port (Fig. 1D). Before assembly, the media inlet, media outlet, and pressure chamber port were punched in the PDMS using a 2 mm-diameter biopsy punch (Militex, Inc.). The measurement capacitor was punched through the upper PDMS slab using a 10 mm-diameter biopsy punch (Accuderm, Inc.), and the reference capacitor was punched with a 5 mm-diameter biopsy punch (Premier Medical Products Company). Templates and guides facilitated pump assembly to standardize fabrication.

**Figure 1.**
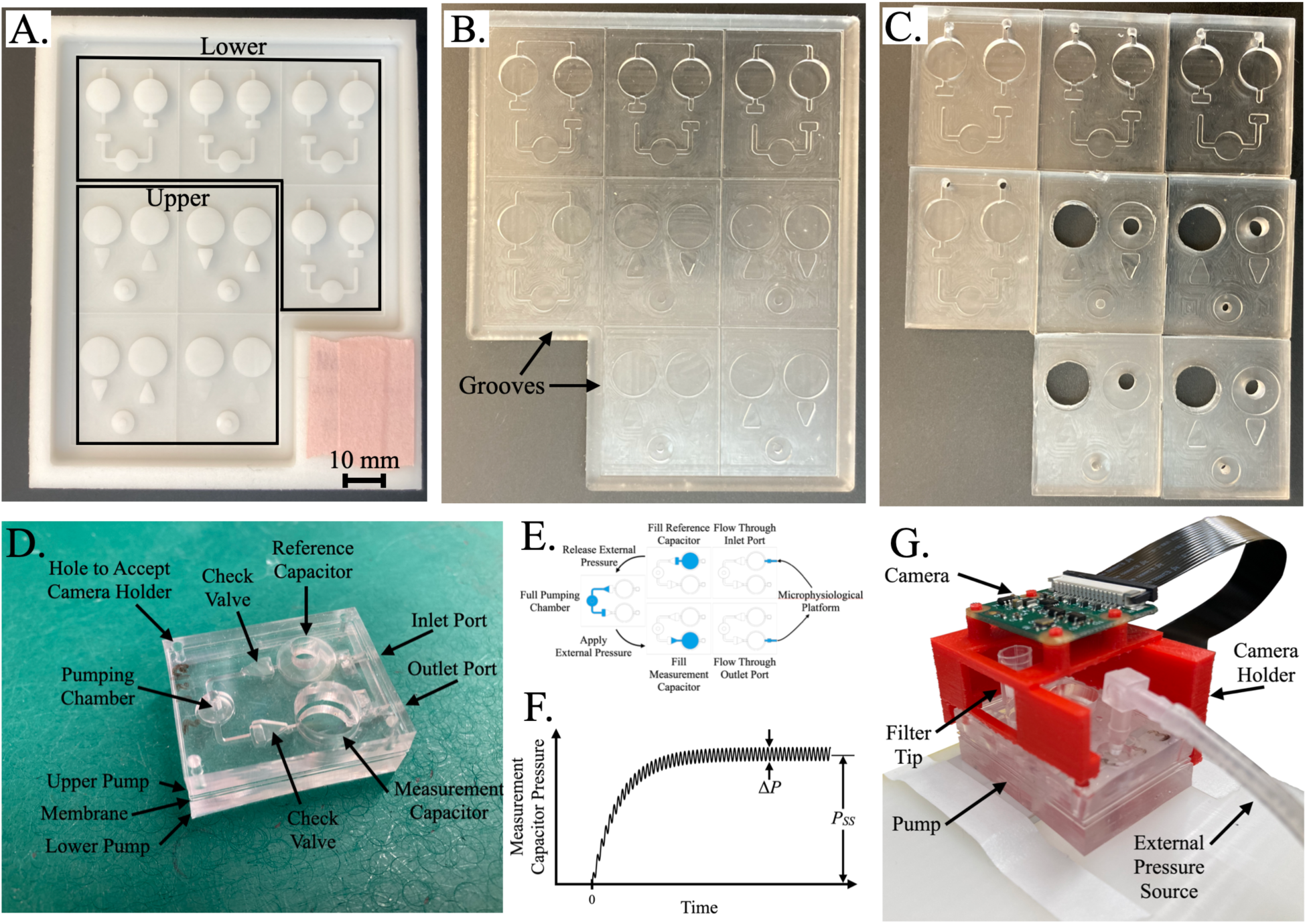
The main components and operation of the pump and optical system. (A) Mold used to cure PDMS to make pump slabs. The mold has features to make four upper and lower pump slabs. Ridges divide the mold into eight individual slabs. (B) Cured PDMS released from the mold with features facing up. The ridges in the mold leave grooves that facilitate accurately separating the cured PDMS into individual slabs. (C) Cured PDMS cut into individual slabs and punched. (D) Assembled pump. The pump consists of a lower slab, silicone membrane, and upper slab. (E) Beginning at the pumping chamber, activating the pump with external air pressure moves fluid through the first check valve and into the measurement capacitor. Fluid is ejected through the outlet port and into the attached microphysiological system or resistive device. Fluid returns through the inlet port and accumulates in the reference capacitor. Releasing the external pressure draws fluid through the second check valve and back into the pumping chamber. (F) Repeated pressurization of the pumping chamber increases the pressure in the measurement capacitor to a steady offset pressure *Pss* with superimposed oscillation of amplitude Δ*P*. (G) A custom-made camera holder was press-fit into holes at the corners of the upper pump. The camera holder had room to insert a filter tip into the reference capacitor and positioned the camera directly over the measurement capacitor.

The key feature of the pumps was their ability to drive flow through an attached device by developing a pressure difference between the measurement and reference capacitors (Fig. 1E). An external pressure source deforms the membrane over the pumping chamber to move fluid into the measurement capacitor (Fig. 1E). Returning the pumping chamber to atmospheric pressure causes valve before the measurement capacitor to close. The pressurized fluid in the measurement capacitor discharges through the microfluidic device and collects in the reference capacitor (Fig. 1E). A typical vascularized MPS offers resistance to flow, so repeatedly pressurizing and releasing the pumping chamber would build pressure in the measurement capacitor pressure to a steady state with small superimposed oscillation (Fig. 1F). Atmospheric pressure was maintained at the air-liquid interface over the reference capacitor by either leaving the port open or covered with a filter tip (Fig. 1G). Four 2 mm-diameter holes were punched in the top of the assembled pump to accept a camera positioning jig (Fig. 1G). The jig accepted a camera module (Raspberry Pi Model 2.1) driven by a single-board computer (Raspberry Pi Model 4). The camera was positioned above the measurement capacitor (Fig. 1G).

### Membrane Preparation

Pressure in the fluid below the membrane was deduced by tracking the motion of fiducial marks affixed to the membrane over the measurement capacitor. Precise positioning of fiducial marks on the membranes was achieved using a series of fixtures and guides (Fig. 2). A 0.3 mm-thick silicone sheet (LMS, Amazon) was cut to a rectangle with approximate dimensions of 4.5 cm by 5.0 cm (Fig. 2A). One guide positioned a 2 mm-diameter punch near the corners of the membrane (Fig. 2B). The holes in the membrane received posts of a fixture (Fig. 2C) that provided alignment between the membrane and guides (Fig. 2C). One guide placed on the same posts positioned a 1 mm-diameter punch over the valves (Fig. 2C and D) [10]. A similar guide positioned an 8 mm-diameter punch (Skler Instruments) over the reference capacitor. A final guide positioned a permanent ink pen (Pilot Corporation of America) to transfer a small amount of ink to the membrane for fiducial marks. The ink pooled and dried into a nearly circular shape (Fig. 2D, inset).

**Figure 2.**
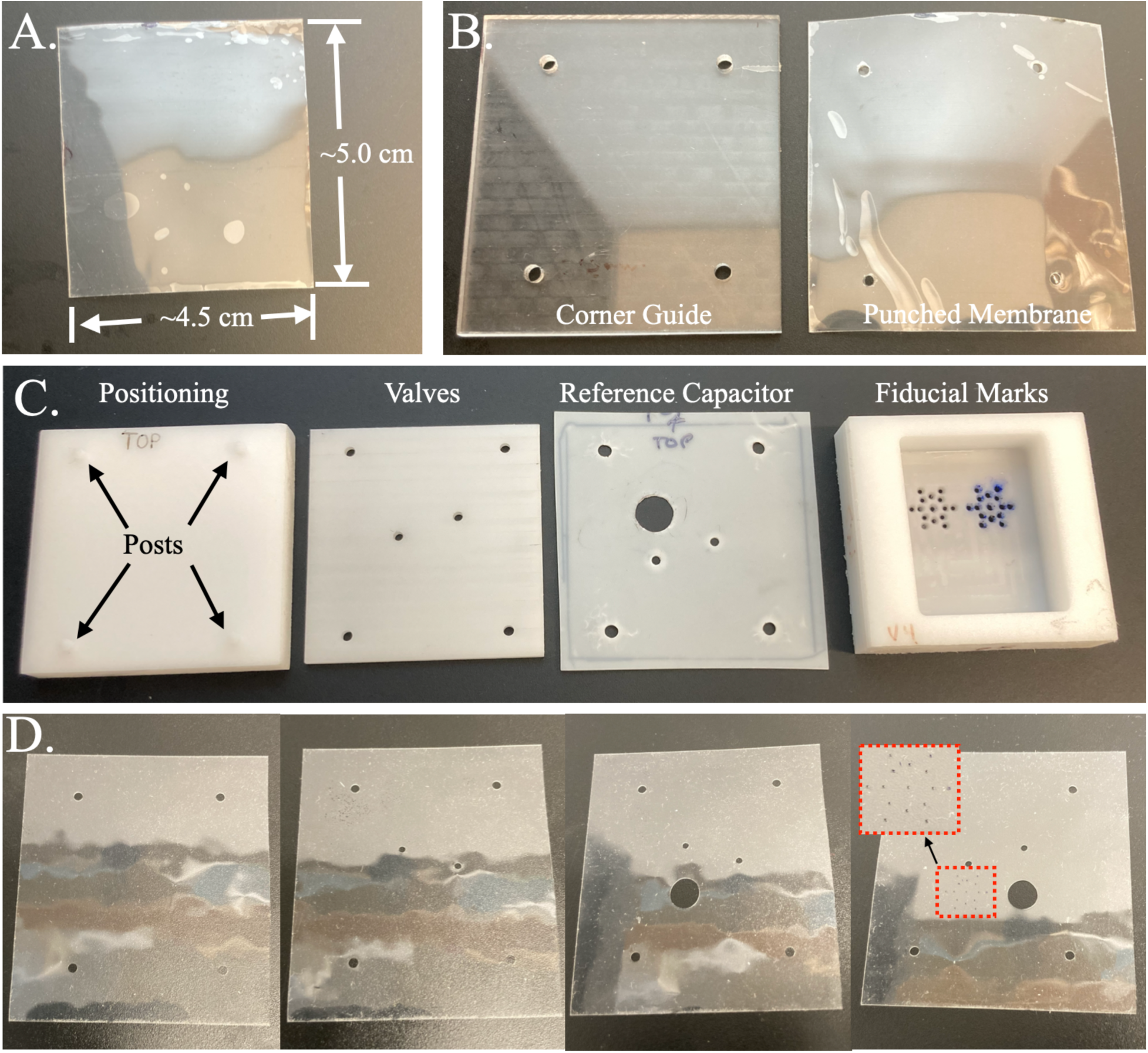
Fixtures and guides to facilitate alignment of the fiducial marks on the deformable membranes. (A) A typical membrane of approximately 4.5 x 5.0 cm before being processed for incorporation into a pump. (B) Acrylic guide used to punch holes in the corners of the membrane and a membrane punched using that guide. The holes in the membrane accept posts of the positioning fixture. (C) The positioning fixture with posts to accept a series of guides to position holes and fiducial marks on the membrane. Various guides to punch holes for the valves and the reference capacitor and to add fiducial marks to the membrane. (D) An unaltered membrane and membranes after punching for the valves, punching for the reference capacitor, and placing the fiducial marks (inset) using the corresponding guide from (C).

### Fiducial Marker Positioning

The fiducial marks served two purposes depending on their position (Fig. 3). “Free” fiducial marks were located on the membrane within the edges of the capacitor and moved as the pressure in the fluid under the membrane changed. “Fixed” fiducial marks were located on the membrane bound to the lower pump slab so their motion was minimal. Fixed fiducial marks served as a reference to correct for spurious motion due to rigid translation and rotation of the membrane relative to the camera.

**Figure 3.**
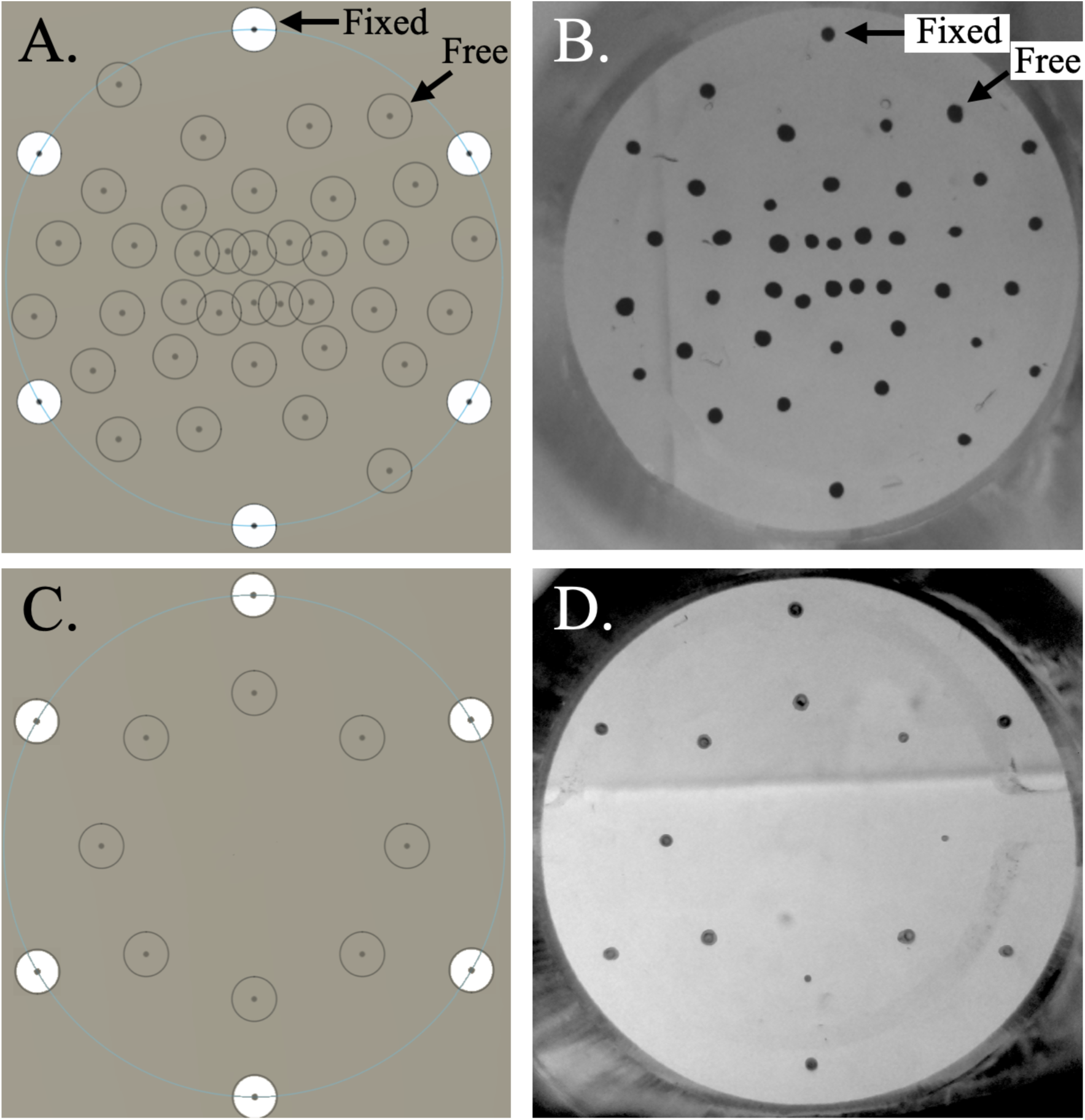
Configuration of fiducial marks on capacitor membranes. (A) Schematic diagram showing positions of fiducial marks at twenty evenly distributed radial positions. The fiducial marks were reflected through the center of the membrane giving a total of 40 fiducial marks. The fiducial marks were staggered circumferentially to facilitate identification and tracking. Fixed fiducial marks shown in white with circumference of the lower capacitor shown by a blue line. (B) Arrangement of fiducial marks on a membrane captured using the optical system. (C) Schematic diagram showing positions of fiducial marks at radial position to maximize radial motion of fiducial marks for a capacitor pressure. (D) Arrangement of fiducial marks on a membrane captured using the optical system. The nearly horizontal line through the membrane appeared at the interface between the two underlying backlit LEDs and was positioned away from fiducial marks.

Two configurations of fixed and free fiducial marks were used to characterize pump function (Fig. 3). The first consisted of an array of 18 free fiducial marks evenly distributed between the center and the edge of the capacitor and staggered circumferentially to facilitate identification and tracking (Fig. 3A). The free fiducial marks were reflected relative to the origin to give a total of 36 free fiducial marks (Fig. 3A and B). Six fixed fiducial marks were uniformly arranged around the circumference of the capacitor.

The second configuration of fiducial marks consisted of eight free fiducial marks with uniform circumferential distribution at a fixed radial position (Fig. 3C and D). Six fixed fiducial marks were uniformly arranged around the circumference of the capacitor (Fig. 3C and D).

### Calibrating Fluid Pressure Under the Measurement Capacitor

Each pump was calibrated to establish the relationship between the measurement capacitor pressure *P*(*t*) and average fiducial marker radial displacement *u*(*t*). Tubing (McMaster-Carr) was filled with fluid and press fit into the pump outlet and connected through a T-fitting to a prefilled horizontal 10 mL syringe (Becton Dickinson), horizontal 3mL syringe (Becton Dickinson), and vertical 5 mL serological pipette (Celltreat) (Fig. 4A). The one-way valve adjacent to the capacitor prevented flow out of the capacitor. The syringes moved fluid into the serological pipette that then acted as a manometer to impose well-defined and easily measured hydrostatic pressure on the measurement capacitor membrane. The capacitor was exposed to six pressures equally distributed between −0.25 to 1.6 kPa, each visited 5 times in a random order, and the position of the fiducial marks under pressure were captured by the imaging system.

**Figure 4.**
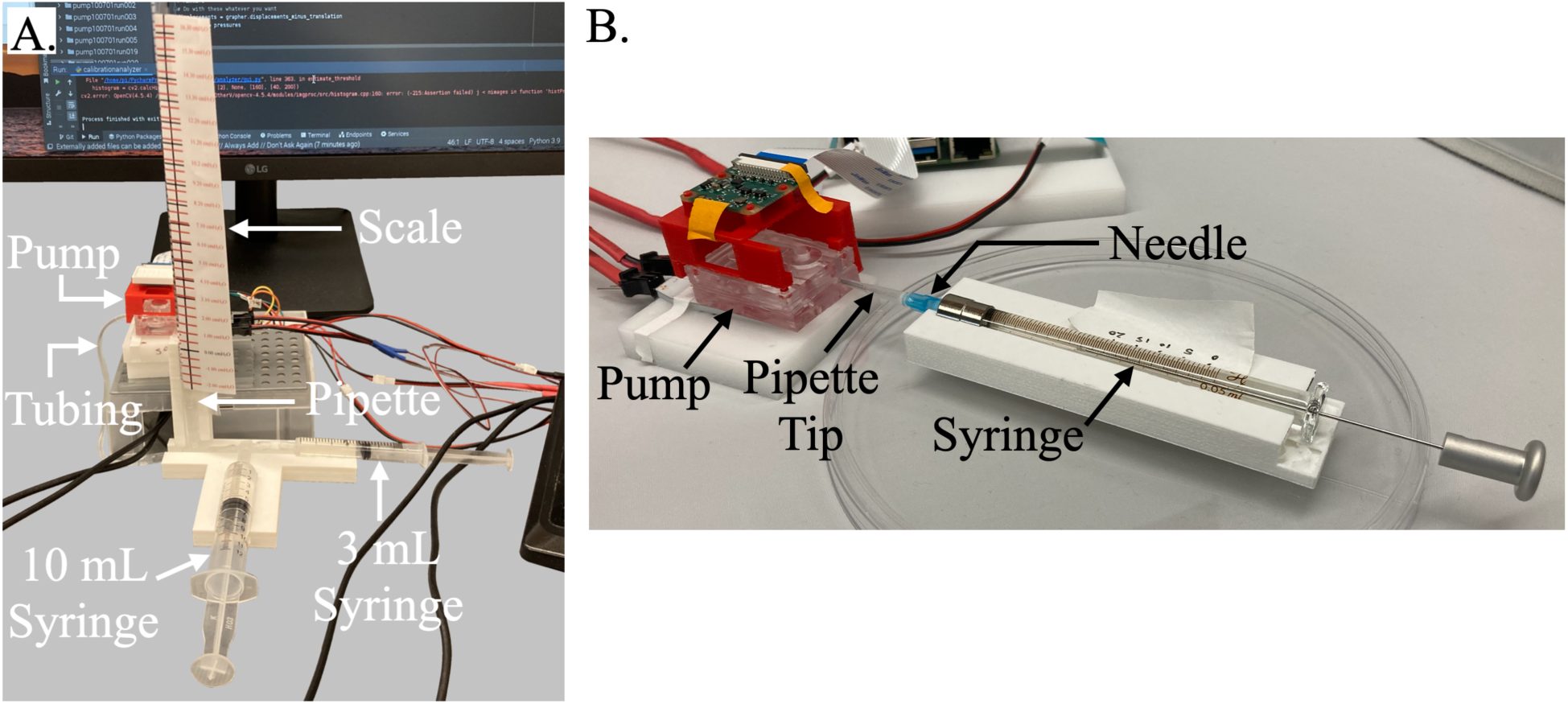
The set-ups used to determine the relationship between measurement capacitor pressure or volume and radial fiducial marker motion. (A) A 10 ml and 3 ml syringe supply fluid to a vertical pipette through a T-fitting. Tubing connects the vertical pipette directly to the measurement capacitor of the pump. The vertical pipette acts as a manometer to apply a well-defined pressure to the measurement capacitor. (B) A 1 ml-syringe was connected to the measurement capacitor through a pipette tip bonding to a needle and press fit into the capacitor outlet.

### Calibrating Added Fluid Volume Under the Measurement Capacitor

The relationship between volume added to the capacitor *V* and the radial fiducial marker displacement *u* was determined using a 1 ml glass syringe (Hamilton) (Fig. 4B). The syringe was connected to the pump through a needle epoxied to a 200 µL pipette tip and press fit into the outlet of the measurement capacitor (Fig. 5B). Known fluid volumes equally distributed between 0 to 20 µL, each visited 5 times in a random order, were imposed on the capacitor, and the fiducial marks on the membrane were captured by the imaging system.

**Figure 5.**
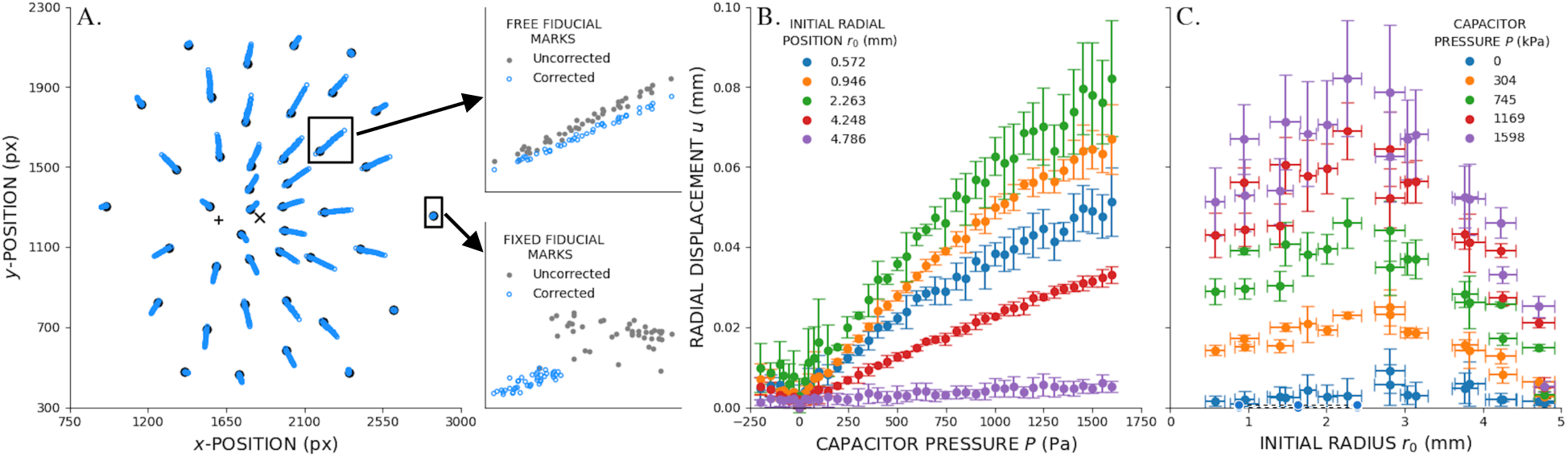
Characterization of fiducial mark motion under known hydrostatic pressures. (A) Initial position of a total of 36 free fiducial marks (black circles) and six fixed fiducial marks and subsequent radial motion under 41 hydrostatic pressures (blue circles). Fiducial mark displacements were scaled 10-fold to facilitate visualization in units of pixels. The motion of a free and fixed fiducial mark shown before and after correcting for spurious motion (insets). Correcting for spurious motion of free fiducial marks caused a small shift from the uncorrected position (gray circles) to the corrected position (blue circles). The position of fixed fiducial marks (gray circles) became more tightly clustered after correcting for spurious motion (blue symbols). The geometric center of the capacitor was indicated with an ✕. The center of motion computed by extending the nearly linear radial trajectories converge at the position indicated with +. (B) Radial displacement of free fiducial marks from several initial radial positions with the capacitor subjected to 41 hydrostatic pressures between −250 and 1600 Pa for a single pump subjected to *n* = 5 calibrations (mean ± standard deviation). (C) Radial displacement of free fiducial marks at five external pressures and 18 initial radii from a single pump subjected to *n* = 5 calibrations (mean ± standard deviation). The maximum displacement occurred at an initial radius around 2.5 mm for all pressures.

### Membrane Capacitance

Capacitance *C* is defined by

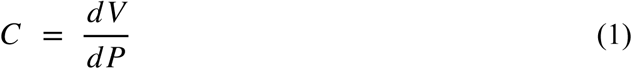

where *dV* is the increment in the volume of fluid contained under the membrane and *dP* is the corresponding increment in pressure of the fluid. The capacitance of the measurement capacitor was approximated using Eq. (1) and the pressure and volume calibrations using

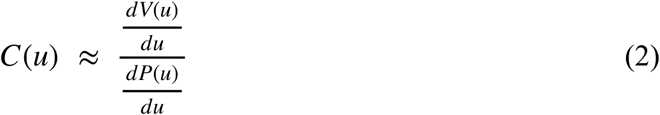

where the numerator *dV*(*u*)/*du* was determined by volume calibration and the denominator *dP*(*u*)/ *du* was determined by pressure calibration.

### Imaging System

Images were captured at a resolution of 3264 × 2464 pixels to ensure each fiducial mark spanned many pixels. Using a common modification to adapt the camera for macroscopic photography, the camera lens was detached from the housing and unscrewed to shorten the focal distance [cf. 11]. The pump and imaging system sat atop a pair of white LED backlight modules (Adafruit Industries) to uniformly illuminate the membrane within the measurement capacitor.

### Image Capture

Image quality was determined in part by parameters set on the camera. These parameters were selected to best identify fiducial marks in the initial configuration based on the ink color and background illumination. The influence of these choices on the performance of the optical system was not investigated systematically. Cursory examination showed that image parameters influenced fiducial marker identification but not tracking.

### Identifying and Tracking Fiducial Marks

The region around each fiducial mark was isolated and the intensity of the blue color channel was subjected to a blurring mask to reduce sensitivity to noise and a threshold map to identify pixels associated with fiducial marks. Pixel intensity below the threshold were set to 0 and greater than or equal to the threshold were set to the maximum pixel intensity of 255. The geometric center of the fiducial mark was computed using a center of mass algorithm [12]. The region around each fiducial mark, the pixels considered part of the fiducial mark, and the position of the fiducial mark were annotated on the captured image. Users were prompted to correct misidentified fiducial marks and confirm fixed and free fiducial marks before initiating the desired protocol.

During a protocol, the computed position of a fiducial mark in the previous image was used to estimate the position of the same fiducial mark in the current image. The time and the computed position of each fiducial mark were saved to a removable SD card. It was possible to also save individual images at frame rate reduced by a factor of ∼5.

### Spurious Motion

The upper pump to which the imaging system was fixed was deformable. The fixed fiducial marks were used to compensate for apparent motion of fiducial marks due to spurious motion of the camera relative to the pump (see Supplementary Material).

### Pump Activation

The pump was driven by an external pressure source that was set by a regulator (Model 700-BA, ControlAir) and controlled through a solenoid (S070C-6CG-32, SMC Pneumatics, [9]). The computer responsible for image acquisition also actuated the solenoid by connecting the pumping chamber to either the regulated pressure or atmospheric pressure.

Pumping was induced by exposing the pumping chamber to a time-dependent external pressure approximated as a square wave. The input waveform was characterized by three parameters: the amplitude, duty cycle, and frequency of the square wave.

### Flow Resistance

The pumps were initially characterized using flow resistance provided by glass capillary tubes with internal radius of ∼70 µm (Hirschmann-Laborgeräte) and lengths that offered flow resistances spanning a range expected of a vascularized MPS. Steady laminar flow at low Reynold’s number through a capillary tube experiences flow resistance *R* approximated by

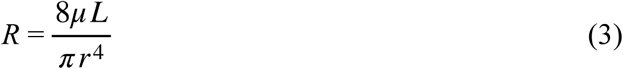

where *µ* is the fluid viscosity, *r* is the internal radius, and *L* is the length of the capillary. Capillary tubes were connected to the pump through the same tubing that fit the outlet port for pressure calibration. The capillary tube was first press fit into a short (∼2 mm) segment of an intermediate sized tubing (McMaster-Carr) filling the gap between the outside of the capillary and the inside of the tubing.

Flow resistance *R* was also determined for a model vascular network composed of branching, interconnected segments. A bifurcating network (Fig. S1) was 3D printed from PLA filament using an extrusion tip of 0.2 mm and a thickness of ∼0.2 mm (Monoprice). The branching network was carefully removed from the build plate and adhered to a petri dish using double-sided tape. PDMS and activator at a ratio of 10:1 was poured over the PLA and cured. The PLA was removed, 2 mm-diameter inlet and outlet ports were punched, and the PDMS with branching features was adhered to a large glass coverslip (Ted Pella, Inc). The branching network was connected to the pump using tubing press fit into the inlet and outlet ports.

### The Effect of Input Pressure Amplitude, Frequency, and Duty Cycle on Pump Performance

The pumping chamber was exposed to seven external pressures *P_EX_* evenly distributed between 0 and 16.5 kPa at constant frequency *f* = 0.5 Hz and duty cycle *D* = 50%. For each cycle of the pressure time course measured at the measurement capacitor, the steady state pressure offset *P_SS_* was calculated as the average pressure and the amplitude of the superimposed pressure oscillation Δ*P* was calculated as the difference between the maximum and minimum pressure of the cycle. The pump was characterized by averaging *P_SS_* and Δ*P* over several cycles. At constant *P_EX_* = ∼8.3 kPa, each of 13 frequencies *f* between 0.01 and 10 Hz and seven duty cycles *D* = 1, 10, 25, 50, 75, 90, and 99% were paired in a random order in a single protocol consisting of 5 min with the pump activated to build to a steady state pressure followed by 5 min with the pump off to return the pressure to baseline. To accommodate the lowest frequency at *f* = 0.01 Hz, the pump was activated for 25 min followed by 5 min with the pump off. The number of cycles included in each calculation of *P_SS_* and Δ*P* was the last nine for *f* = 0.01 and 0.03125 Hz and the last ten cycles for *f* > 0.03125 Hz.

### Quantifying Flow Resistance by Hydrostatic Pressure

Flow resistance *R* of each capillary tube with associated tubing was quantified by measuring the mass flow rate driven by a constant hydrostatic pressure Δ*P*. A 60-mm diameter petri dish filled with water was connected to a capillary tube that discharged into a second petri dish on a scale (Mettler-Toledo). The 60-mm diameter petri dish was elevated to induce flow and the mass of water deposited on the scale was recorded at discrete times, *m*(*t*). Volume flow rate was computed by dividing the mass flow rate by the density of water, 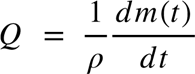. Flow resistance *R* was estimated using

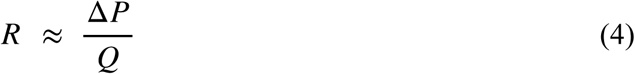

### Quantifying Flow Resistance by Pressure Decay

Quantifying flow resistance using the optical system involved activating a pump at a frequency of 0.5 Hz and duty cycle of 50%. The system ran for 3 min to allow the pressure in the measurement capacitor *P*(*t*) to reach a steady state. The optical system measured *P*(*t*) for 20 s to quantify *P_SS_* before the pumping chamber was connected to atmospheric pressure. Capacitor pressure *P*(*t*) decreased as fluid moved from the pressurized measurement capacitor, through the attached capillary, and into the reference capacitor. When *P*(*t*) dropped below two-thirds of *P_SS_*, the pump was reactivated for 3 min and the protocol was repeated. Given the constant capacitance assumption, it was expected that *P*(*t*) would decay exponentially according to

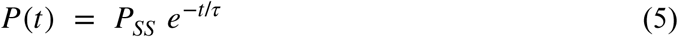

where *P_SS_* is the steady-state pressure at the time the pumping chamber was connected to atmospheric pressure, *t* is time, and τ is a time constant characterizing the rate of pressure decay due to flow out of the capacitor. The time course of *P*(*t*) allowed computation of *P_SS_* from the average pressure measured in the 20 s prior to the arrest of pumping, and τ was estimated from a linear fit of ln *P*(*t*) vs. *t*. By analogy to flow of electric current through a resistor and capacitor in series, = *R* ⋅ *C* and the resistance of the attached device was estimated from

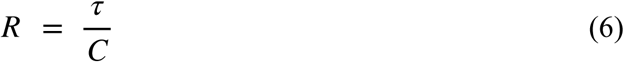

### Resistance of Microvascular Networks

Vascular networks in a microphysiological device were created as described previously [13]. Human umbilical vein endothelial cells (ECs) were grown in tissue culture flasks in Vasculife (Lifeline Cell Technology). Normal human lung fibroblasts (FBs) were grown in Fibrolife (Lifeline Cell Technology). The cells were detached and resuspended in a solution of media and fibrin. The cell slurry was mixed with an equal volume of thrombin at 2 U/ml and injected into the microphysiological platform at a final concentration of 6.0×10^6^ endothelial cells, 1.5×10^6^ fibroblast cells, and 3.0 mg/mL fibrin. The system was maintained under static conditions in a humidified chamber in an incubator for 5 days before being connected to a pump. The optical system determined *R* of the vascular network every 2 h over ∼48 h. Immediately before connecting the pump to the device and immediately following the experiment, fluorescent dextran was flown through the device to confirm network perfusability.

## Results

### Pressure and Volume Calibration

Each pump was filled and fiducial marks were captured when the pressure in the measurement capacitor was atmospheric, *P* = 0. Changing *P* within the measurement capacitor between −250 and 1,600 Pa resulted in membrane deformation detected by the optical system as radial motion of the fiducial marks (Fig. 5A). The outermost array of fixed fiducial marks showed little motion relative to their position in the initial image (Fig. 5A, inset). The parameters that accounted for spurious translation, rotation, and dilation from all 5 pumps used for calibration were Δ*x* < 10 pixels, Δ*y* < 5 pixels, θ 0, and λ 1 (Fig. S2). These parameters resulted in very small corrections to the absolute positions of the free fiducial marks (Fig. 6A). Although small, correcting for spurious motion tightened the clustering of fixed fiducial marks (Fig. 6A, insets). Over the full range of capacitor pressures, the corrected motion of the free fiducial marks was remarkably radial (Fig. 6A). Projecting the radial motion of the free fiducial marks to the center of the capacitor showed that the center of motion of the fiducial marks was not always at the geometric center of the capacitor (Fig. 6A). Nevertheless, the average radial displacement *u* increased nearly linearly with reference capacitor pressure *P*(*t*) (Fig. 6B). It is noteworthy that *u* also depended on the initial radial position of the fiducial mark *r*_0_ (Fig. 6B). Radial displacement at a given capacitor pressure *P* increased to a maximum and then decreased with increasing *r*_0_ (Fig. 6B and C). Measuring *u*(*r*) over several membranes revealed that the maximum *u*(*r*) at each *P* occurred at or slightly above the radial position *r_C_* 2.5 mm (Fig. 6C). Membranes with fiducial marks at *r_C_* is desirable because it maximizes the motion of fiducial marks for a given *P*, increasing the sensitivity of the pressure or volume measurement. The second membrane configuration included all free fiducial marks positioned at *r ≈ r_C_* (Fig. 3 C and D).

**Figure 6.**
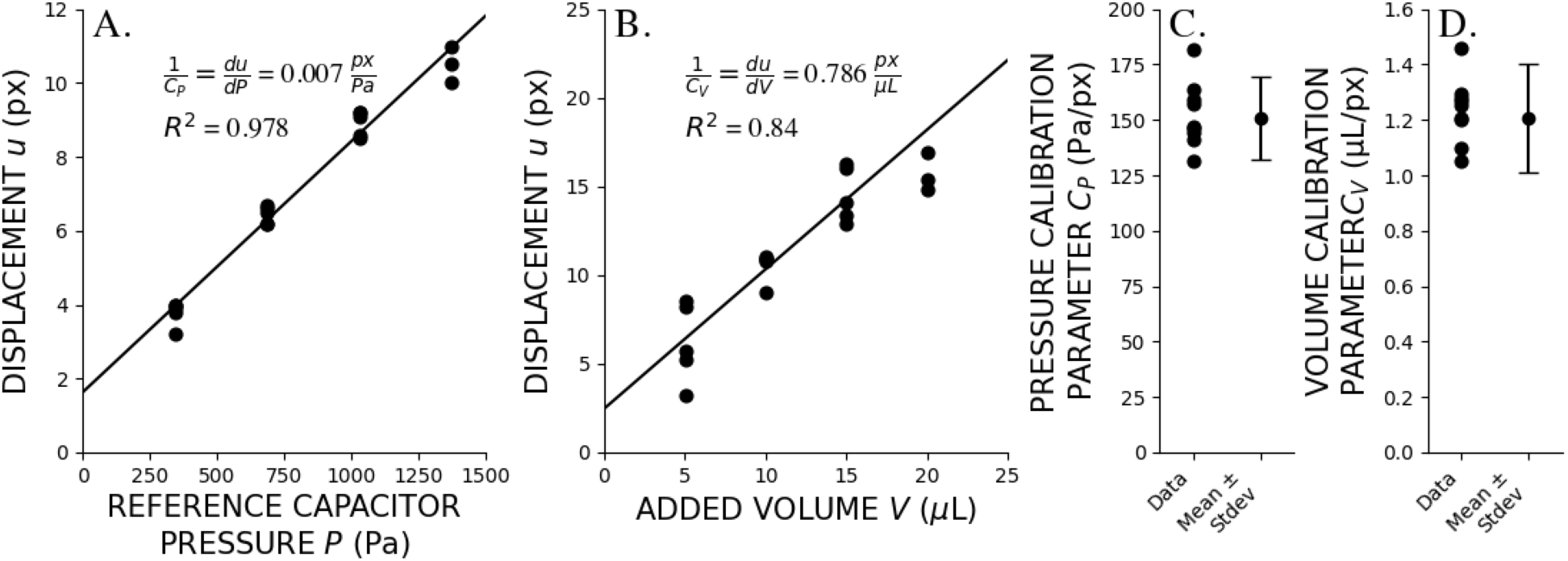
Calibration parameters. (A) Representative calibration data showing capacitor radial displacement for hydrostatic pressures up to ∼1,300 Pa with each pressure visited five times in a random order. Superimposed linear regression showing the slope 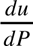 and coefficient of correlation *R*^2^. (B) Representative calibration showing capacitor radial displacement for added volumes up to 20 µL with each volume visited five times in a random order. Superimposed linear regression showing the slope 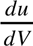 and coefficient of correlation *R*^2^. (C) Pressure calibration parameter data for all fourteen pumps used in this study (mean ± standard deviation). (D) Volume calibration parameter data for all fourteen pumps used in this study (mean ± standard deviation).

Pumps with free fiducial marks at *r_C_* were individually calibrated by applying known hydrostatic pressures and infusing known fluid volumes directly into the measurement capacitor. Radial displacement of the free fiducial marks *u*(*r*) increased nearly linearly with either *P* or Δ*V* (Fig. 6A and B). Thus, each pump was completely characterized by the slopes of the calibration curves 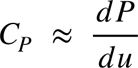 and 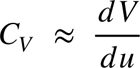. Taken together, the calibration parameters for all pumps were similar, giving mean and standard deviation of *C_P_* ≈ 151.0 ± 18.5 Pa/pixel and *C_V_* ≈ 1.21 ± 0.19 µL/pixel for *n* = 14 pumps (Fig. 6C and D).

#### Dynamic Performance

The system captured an image, identified fiducial marks, determined marker position, computed marker motion, and estimated capacitor pressure at a rate of about 14 Hz. With no external air pressure driving the pump, the average motion of the free fiducial marks was 0.143±0.028 pixels, corresponding to a level of noise in apparent horizontal displacement, vertical displacement, capacitor pressure, and capacitor volume of 0.714±0.14 µm, 0.827±0.16 µm, 22.5±4.5 Pa, and 0.180±0.036 µL, respectively. The pump was connected to a capillary with *L* = 32 mm and activated by *P_EX_* = ∼8.3 kPa at *f* = 0.5 Hz and *D* = 50% (*i*.*e*., on and off times of 1 s). Fluid accumulated in the measurement capacitor causing *P*(*t*) to increase to a nearly steady state value *P_SS_* of ∼250 Pa (Fig. 7A). The steady state pressure was reached after ∼60 s and maintained for ∼4 min (Fig. 7A). On a shorter timescale, the system captured the small variations in capacitor pressure corresponding to the 0.5 Hz pressurization and venting of the pumping chamber (Fig. 7A and insets). Thus, the optical system demonstrated that pumps drive flow through an attached resistance by generating and maintaining a nearly steady fluid pressure with small superimposed oscillation.

**Figure 7.**
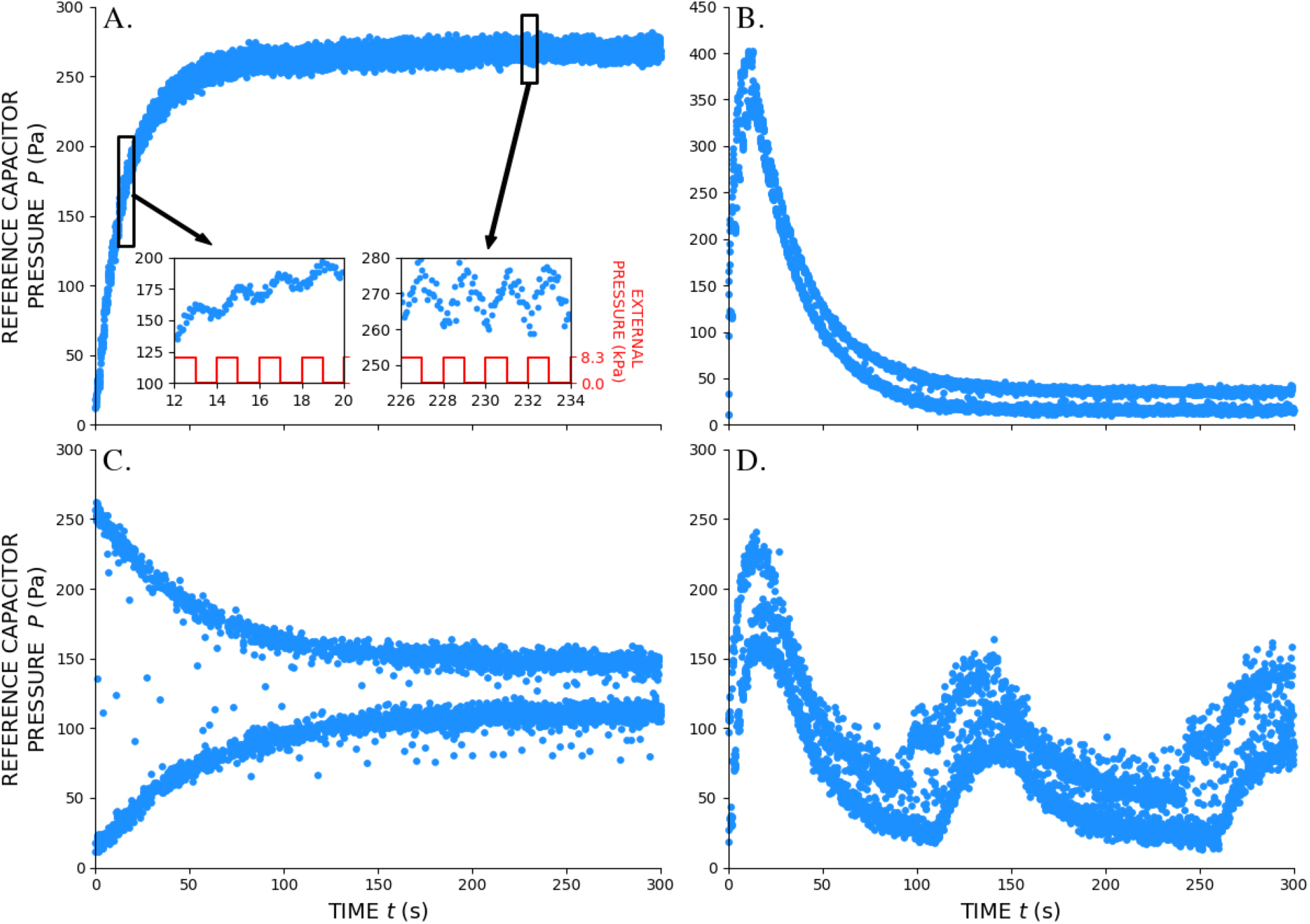
Representative pump responses. (A) Pressure in the measurement capacitor steadily increased to reach a near steady state pressure. The optical system detected the superimposed oscillations with the external pressure applied at the pumping chamber both during the build-up of pressure and at steady state (Insets). (B) A pump that supported an increase in pressure then experienced decrease in pressure despite the input pressure waveform continuously imposed on the pumping chamber. The system detected oscillation due to the input pressure. (C) The pressure in the measurement chamber oscillates between a high-pressure curve and low-pressure curve that appear to be converging around 125 Pa. The system detects very few instances where the capacitor is between these extremes suggesting that the membrane is moving almost instantaneously with the pressure supplied to the pumping chamber. (D) A pump exposed to the same input waveform but exhibited behavior that is not easily explained. Only pumps that qualitatively behaved like the pump displayed in (A) were used in this study.

The optical system identified pumps exhibiting poor or unpredictable performance. Some pumps exhibited a spontaneous decrease in pressure despite continuous external pressurization of the pumping chamber (Fig. 7B). Curiously, the capacitor pressure experienced detectable fluctuations suggesting fluid continued to be pumped into the measurement capacitor as *P*(*t*) decayed. Other pumps showed large oscillations in *P* with pumping (Fig 7C). The imaging system detected the membrane almost exclusively at two configurations corresponding to the pumping chamber pressure *P_EX_* = 0 or ∼8.3 kPa. The behavior of some pumps was not easily explained (Fig. 7D). Interestingly, simply determining if a pump moved fluid would be woefully inadequate in identifying poorly performing pumps since all pumps in Fig. 7 move fluid. Only pumps in which the optical system detected an increase in capacitor pressure to a steady value during pumping (*e*.*g*., Fig. 7A) were used in subsequent experiments. A minority of manufactured pumps exhibited the behavior shown in Fig. 7A.

The magnitude of *P_EX_* strongly influenced *P_SS_* and weakly influenced Δ*P* (Fig. 8A). Capacitor pressure increased rapidly upon initiating pumping with the time required to reach *P_SS_* about 60 s for all *P_EX_* (Fig. 8A). The rate of *P*(*t*) decay was slower taking ∼120 s to return to baseline (Fig. 8A). Both *P_SS_* and Δ*P* increased nearly linearly with *P_EX_* between 6 and ∼17 kPa with *P_SS_* increasing more rapidly than Δ*P* (Fig. 8B). Removal of *P_EX_* caused the *P*(*t*) to decrease as the capacitor membrane recovered its undeformed configuration and fluid was ejected through the outlet port. The logarithm of *P*(*t*) decreased nearly linearly for all *P_EX_* > 0 over the first 60 s after ceasing pumping (Fig. 8C, inset). Over all examined *P_EX_*, the logarithm pressure decreased nearly linearly with time following Eq. (5) with a minimum *R*^2^ of 0.994. The rate of pressure decay was used to assess flow resistance using Eq. (6).

**Figure 8.**
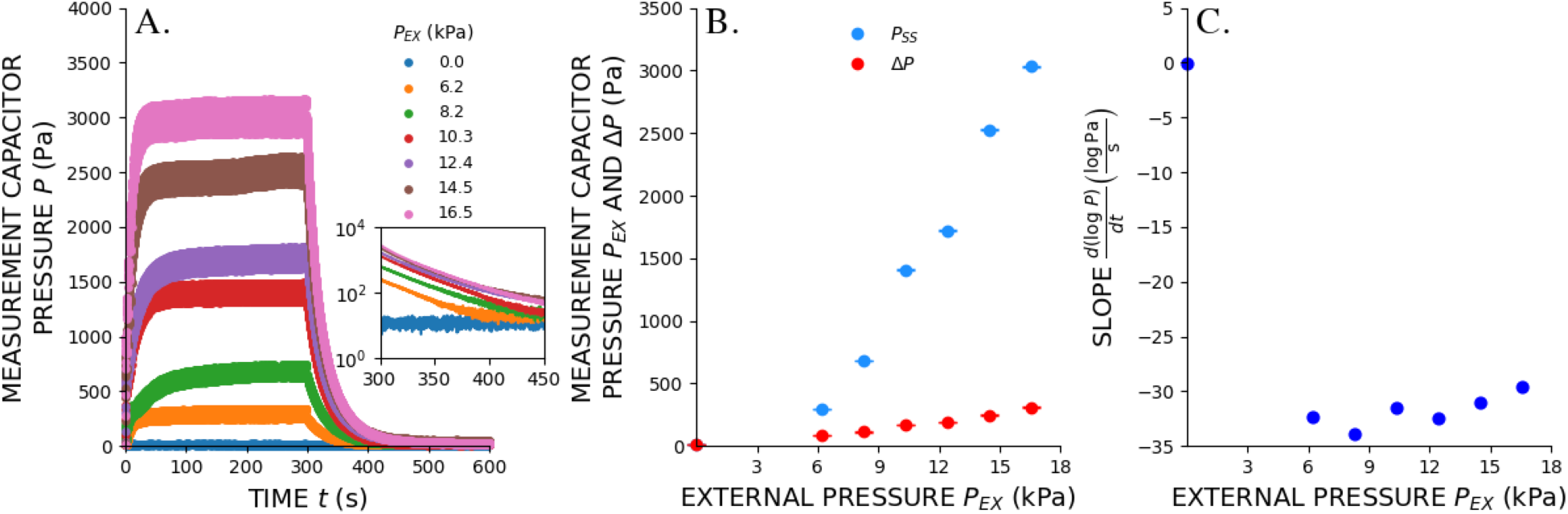
The role of external pressure on the steady-state and superimposed oscillation pressure measured at the measurement capacitor. (A) Time course of capacitor pressure at various external pressures imposed on the pumping chamber. The solenoid switched the pumping chamber between the external pressure and atmospheric pressure for the first 300 s before opening to atmosphere for the remaining 300 s. The initial decay of the logarithm of pressure was nearly linear (inset). (B) The steady state *P_SS_* and superimposed oscillation pressure Δ*P* determined during the 10 second before ceasing pumping (mean ± standard deviation from *n* = 10 cycles). (C) The slope of the logarithm of pressure as a function of time for various external pressures.

The influence of input frequency *f* and duty cycle *D* at constant *P_EX_* on *P_SS_* and Δ*P* was examined as potential means to control pump behavior through only activation of the solenoid (Fig. 9). For *P_EX_* = ∼8.3 kPa and flow resistance provided by a capillary tube with *L* = 32 mm, *P_SS_* for duty cycles of 1 and 99% was low with little frequency dependence (Fig. 9A). At intermediate duty cycles, *P_SS_* increased systematically with *f*. This suggests that the measuring capacitor cannot build pressure when the solenoid spends most of the period connecting the pumping chamber to regulator pressure or atmospheric pressure. *P_SS_* reached a maximum at *D* = 50 (Fig. 9B). Pressure oscillations Δ*P* decreased with *f* (Fig. 9C). At low *f*, Δ*P* increased slightly before decreasing with *D* (Fig. 9D). At higher frequencies (*f* ≥ ∼1 Hz), Δ*P* showed little dependence on *f* or *D* (Fig. 9D).

**Figure 9.**
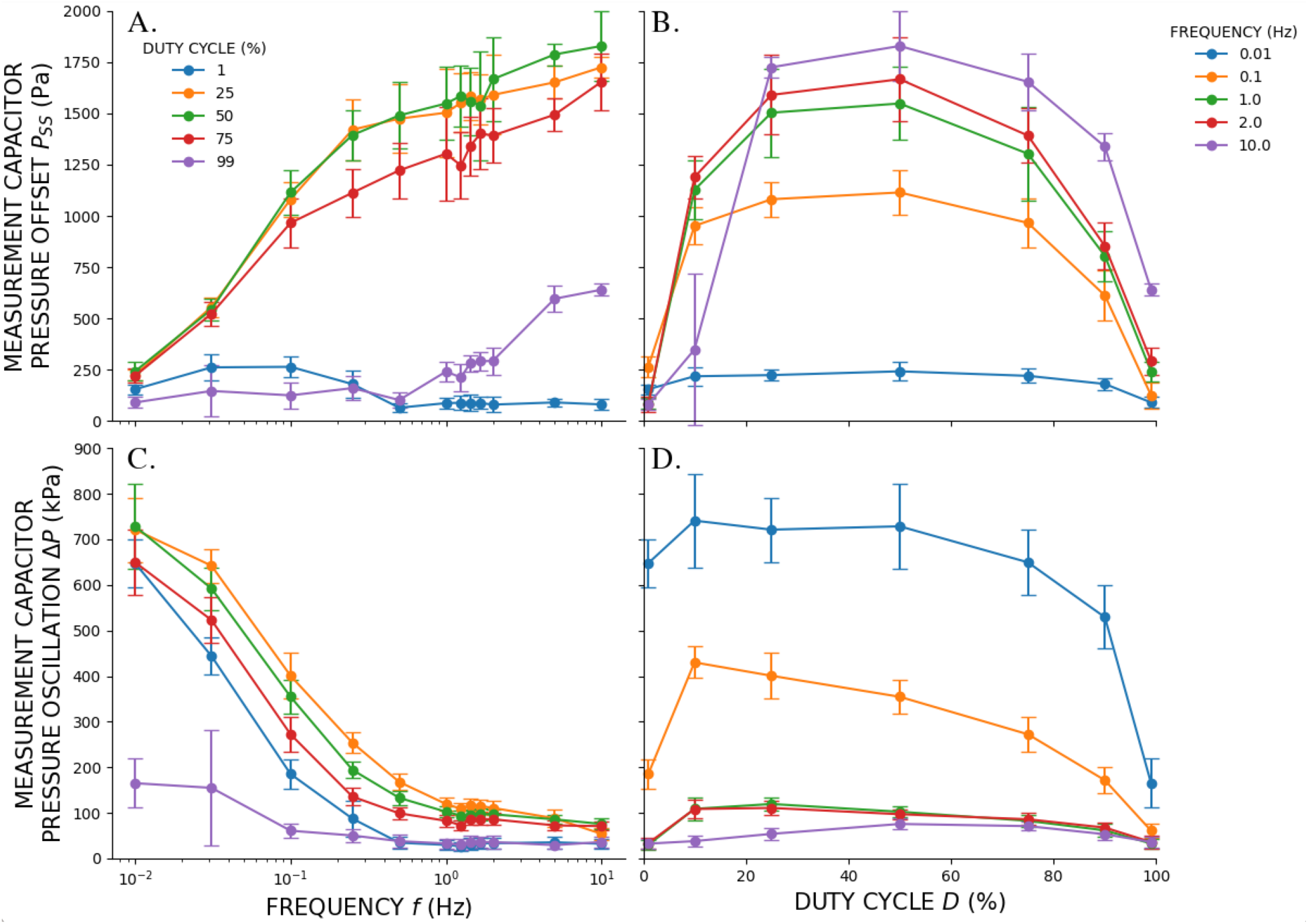
Effect of duty cycle and frequency on pressure offset and oscillation for an input waveform with period of 2 s supplied to the pumping chamber. Data are mean ± standard deviation averaged over 32 ≤ *n* ≤ 90 cycles depending on the frequency. (A) Pressure offset as a function of frequency for several duty cycles. (B) Pressure offset as a function of duty cycle at several frequencies. (C) Pressure oscillation as a function of frequency for several duty cycles. (D) Pressure oscillation as a function of duty cycle at several frequencies.

Flow resistance of glass capillary tubes was determined by measuring the mass flow rate under constant hydrostatic pressure. The mass of water that flowed through the capillary tube increased nearly linearly with time (Fig. 10A). Reynold’s number of the flow within the capillary tube was computed using 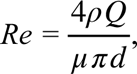 where *ρ* = 0.997 g/mL is the density, *µ* = 1.002 mPa·s was the kinematic viscosity of water at room temperature, *d* 140 µm was the internal diameter of the capillary, and *Q* was the volume flow rate. The range of *Q* measured here gave 0.3 ≤ *Re* ≤ 3. Finding *Re* several orders of magnitude below the critical value 2,300 associated with the transition from laminar to turbulent flow in a tube justified assumption of laminar flow inherent in Eq. (3). The flow resistance for capillary tubes ranging in length from *L* = 0 mm (*i*.*e*., just the associated tubing) to 64 mm increased nearly linearly with capillary length (Fig. 10B). Except with the longest capillary length *L* = 64 mm, the flow resistance was qualitatively similar but slightly higher than the theoretical resistance for laminar flow through a smooth channel (Fig. 10B). Resistance of the same capillaries and tubing was also measured using the optical system. The logarithm of pressure decreased nearly linearly with time (Fig. 10C, inset). The optical system quantitatively and qualitatively captured the dependence of flow resistance on capillary length measured by the hydrostatic method and predicted by theory. Measurements between the optical and hydrostatic methods only differed at *L* = 64 mm. Interestingly, the resistance determined by the optical system for the highest resistance capillary at *L* = 64 mm was below the value computed by hydrostatic pressure but extremely close to the value expected for laminar flow through a smooth tube (Fig. 10B).

**Figure 10.**
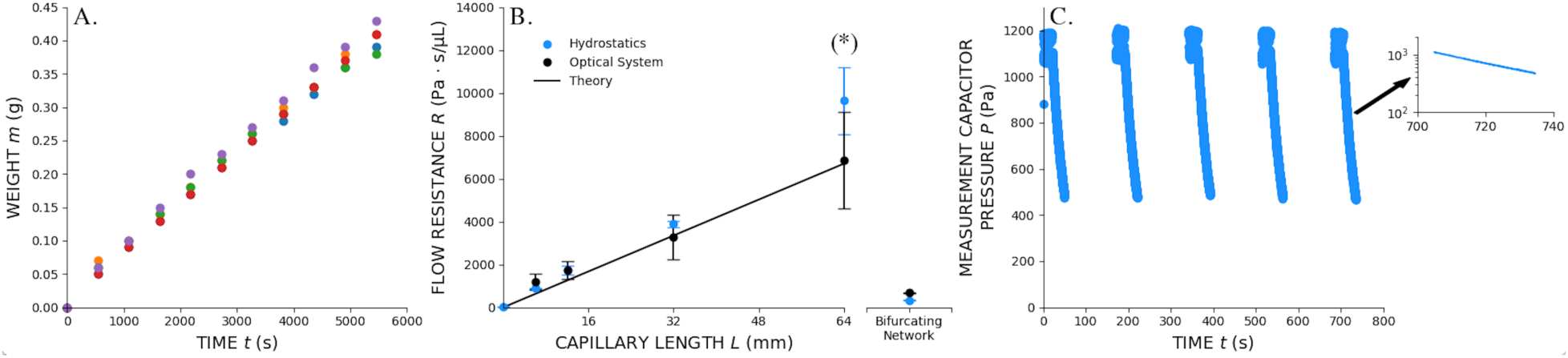
Measurements of flow resistance using the decay in capacitor pressure. (A) Measurement of the mass of water discharged from a capillary tube with length *L* = 32 mm subjected to a hydrostatic pressure of 300 Pa. Independent experiments shown in different colors. (B) Resistance of capillary tubes of various lengths *L* determined by the hydrostatic (blue circles) and optical method (gray circles) (mean±standard deviation for *n* = 5). Each symbol represents an independent experiment. The resistance of a smooth cylinder subjected to laminar flow is superimposed over the experimental data (black line). The flow resistance of a model capillary network determined by the optical and hydrostatic methods. (C) Decay in capacitor pressure determined by the optical method for a pump connected to a capillary tube with *L* = 32 mm. The decay in logarithm of pressure is a nearly linear function of time (inset).

The resistance of the branching network determined by hydrostatic pressure was lower than the smallest capillary (Fig. 10B). The values measured by the optical system were similarly small but overestimated the resistance of the model microvascular network (Fig. 10B). Taken together, the optical system displayed strong quantitative agreement with measurements of flow resistance measured independently and predicted by theory.

Having established that the optical system is able to quantitatively determine the driving fluid pressure of the pump and the flow resistance of the attached device, the optical system was attached to a pump driving flow through a microvascular network growing in an incubator over ∼48 h with *P_SS_* and *R* measured every two hours (Fig. 11). *P_SS_* increased during the four hours immediately after connecting the pump to the microphysiological device and exposing the vascular network to flow (Fig. 11A). It then oscillated around a nearly constant offset of *P_SS_* 1,500 Pa with Δ*P* = 200 Pa over the next ∼24 h. The steady state pressure offset *P_SS_* increased to ∼2,200 Pa around the time when the media in the pump and MPS was changed (Fig. 11A). Between resistance measurements between *t* = 26 and 28 h, the pump was disconnected from the MPS, air bubbles within the pump were noted, and the media was changed. After reconnecting the optical system and reinitiating pumping, capacitor pressure recovered at approximately the same rate as was seen upon the initiation of flow. The first resistance measurement was ∼24 kPa·s/µL corresponding to the published resistance of a microvascular network that was grown for 5 days under static conditions and only exposed to intravascular flow long enough to make the resistance measurement (Fig. 11A, inset; [14]). The final resistance measurement after ∼48 h of flow was higher at ∼32 kPa·s/µL (Fig. 11A, inset). The latter value was measured after disconnecting the pump from the microphysiological device, changing the media, and resetting the reference configuration of the fiducial marks. Some intermediate times gave unrealistically large or small resistance values. However, the initial, final, and several intermediate values obtained using the optical system were quantitatively similar to the flow resistance measured by a different hydrostatic method applied to vascular networks grown in the same lab but tested outside an incubator (Fig. 11B; [14]).

**Figure 11.**
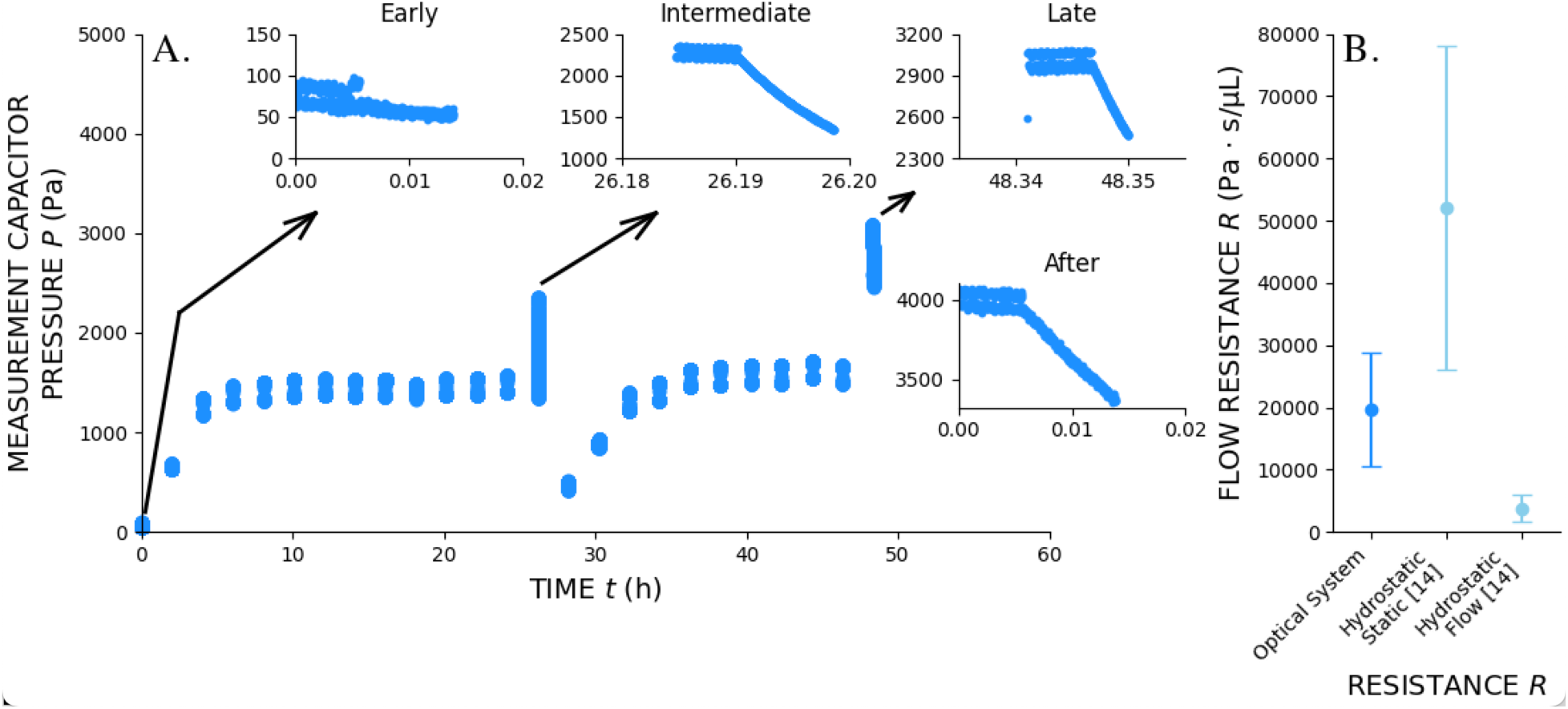
Measurements from a microphysiological platform supporting a vascular network within an incubator. (A) Time course of capacitor pressure measured every 2 h for ∼50 h. The response of the device from an early, intermediate, and late timepoints showing the mean capacitor pressure assessed over 20 s before the pumping chamber was exposed to atmospheric pressure to observe pressure decay. Pressure decay measured outside the incubator after 50 h in the incubator. (B) Comparison of resistance of vascular networks obtained before, during, and after initiating flow using the optical system in the current study (*n* = 4) and vascular networks grown under static or flow conditions using a hydrostatic approach in [14].

## Discussion

This study described a novel and non-invasive optical method for quantifying key parameters determining flow of media through an avascular gel or microvascular network within a MPS. This represents significant advancement in the overarching goal of developing a cost effective and approachable method to quantify the flow properties delivered by a pump to a MPS supporting long-term development of vascular structures.

Fluid flow regulates cell biology and emergent behaviors in MPS. Interstitial flow acutely enhances network perfusability through changes in vessel morphology [14-18] that seems to originate in vascular endothelial cells [14]. Sustained flow alters barrier function [16, 18] and the inflammatory state of endothelium [16, 18]. Interestingly, flow in MPS also influences the behavior or circulating cells. Cancer cells introduced into a MPS and exposed to intravascular and interstitial flow exhibit enhanced motility while intravascular and extravascular [19]. Phenotypic changes occur rapidly and may manifest at the level of genes or proteins [18, 20]. The growing body of evidence indicating sensitivity of vascular cells, supporting cells, and circulating cells to physiological flows underpins the importance of controlling environmental forces to recapitulate physiological function in MPS.

The optical system was developed for a pump specifically designed to interface with MPS [9]. Other approaches to subject MPS to flow including hydrostatic reservoirs [2, 15-17, 19-21], syringe pumps [cf. 22, 23], and peristaltic pumps [22, 24, 25]. However, the pump was particularly attractive because maintained a constant driving pressure, allowed recirculation of media, and required only a small media volume. The pumps are likely to be available to any lab already working with PDMS-based MPS since their manufacture uses the same techniques and equipment.

Monitoring flow resistance of MPS over time provides insight into aspects of vascular development and remodeling [26] that are typically only available through microscopic techniques. Measured flow resistances were extremely close to the values predicted for laminar flow through a smooth capillary and were in good agreement with the values computed using an independent experimental procedure based on hydrostatically driven flow. The largest discrepancy between the hydrostatic method and the optical method was observed for the highest resistance where two capillary tubes connected in series to give a total capillary length of 64 mm. The measurement of the optical system was so close to theoretical prediction that it raises the possibility that the hydrodynamic method becomes inaccurate at high resistance. After testing the system with capillary tubes and a model vascular network, the optical system measured flow resistance of an intact vascular network in an incubator over ∼48 hours, with some caveats. Some measurements showed little decay in pressure after stopping pumping, a phenomenon consistent with a bubble impeding flow. Other measurements of vascular resistance were in good agreement with published results [14].

The value of the system might extend beyond quantitative values of vascular resistance to include changes in vascular resistance over time. For example, resistance to flow would be very high before vasculature forms since the driving pressure at the pump can only force flow through the dense fibrous network (hydrogel) occupying the interstitial space. Resistance would decrease as vascular formation proceeded and an alternative route for flow developed. Vascular remodeling is induced by intravascular flow [18]. Measuring vascular resistance in real-time would reveal perfusability without the burden of microscopic observation of vascular structures. In studies where labeling endothelial cells to image vascular structures is undesirable or impossible, this system would provide an alternative approach to infer network perfusability.

To study the role of fluid forces on physiological function in MPS, it might be desirable to maintain a constant pressure drop across the MPS as vascular networks evolved. This could be accomplished by manually adjusting the regulated air pressure supplied to the pumping chamber in real-time. Another approach more amenable to control using the optical system would be to adjust the pressure drop using only the solenoid. Small changes in the frequency and duty cycle of the external driving pressure waveform produced continuous changes in *P_SS_* and Δ*P* (Fig. 9). This revealed an attractive feature of the optical system that could be used to control the properties of the pump without adjusting the air pressure at the regulator. In a study where multiple devices are being exposed to flow simultaneously, an alternative to having dedicated pressure regulators and solenoids for each pump is to have all pumps connected to a single pressure regulator and control pump behavior using the optical system and multiple inexpensive solenoids, each under independent control. The pump could be programmed to maintain a constant driving pressure or steady flow rate as each vascular network evolves over time. Similarly, MPS could be exposed to constant flow rates by measuring *P_SS_* and *R* and using Eq. (4).

The initial pump design featured an intact membrane over the reference capacitor [9]. The first iteration of the optical system included two cameras to simultaneously monitor the pressure in both the measurement and reference capacitor. While this configuration could be appealing in some contexts, opening the reference capacitor membrane simplified the approach by requiring only one camera to assess the pressure driving flow through the attached device. In addition to providing a well-defined reference pressure, the open capacitor provides an opportunity to sample or change media.

The optical system showed that pump behavior can be surprisingly unpredictable (Fig. 7). Simply observing that a pump is capable of moving fluid would be inadequate to identify properly functioning pumps since all pumps generating positive pressure induce flow. Unpredictable pump behavior would result in vasculature being perfused by unexpectedly low or highly variable driving pressures (Fig. 7C and D). One possible source of unpredictability is the function of the valves, although additional investigation outside the scope of this paper did not reveal an obvious underlying cause.

The location of the fiducial marks on the membrane that maximizes the radial displacement under pressure was based on experimental observations obtained by calibration and supported by engineering models of thin, elastic membranes undergoing large deformation ([27]; Supplementary Material). The model predicted a maximal radial displacement for the region of the membrane slightly outside the radial midpoint. This observation was a key element in the design of the current system. Optimizing radial displacement at a fixed capacitor pressure assumed the fiducial marks were arranged symmetrically about the geometric center of the capacitor. Alignment was the goal but was not assured by the manufacturing process. Minor misalignment of the fiducial marks and lower pump had little impact on the ability of the pump to quantify capacitor pressure since each pump was individually calibrated. However, optimizing the system will require minimizing or eliminating this misalignment. One approach to improve pump fidelity is to automate some of the manual steps in marking, punching, and assembling the pumps. Reducing the number of guides and fixtures by incorporating features into the mold to eliminate punching or assembling pumps in bulk instead of individually might reduce the number of steps where assembly could introduce misalignment. Another approach to improve the optical system is to accept some misalignment in the manufacturing process and account for it through the fiducial marker tracking software. Models of membrane mechanics have been established based on very reasonable assumptions - the membrane is thin, elastic, and subjected to finite deformation. The experimental observations are in very good agreement with the model predications (Supplementary Material). Leveraging engineering models is appealing because it may not be possible to eliminate the misalignment, but it would be possible to use imperfect fiducial marker positioning to admit an accurate inference of membrane motion. Having tighter correspondence between the motion of the fiducial marks and the membrane motion could be useful if the optical system is incorporated in a feedback system designed to maintain a specific pressure or flow rate.

Incorporating the optical system into a pump connected to a MPS supporting a vascular network revealed some shortcomings that need to be addressed for the difficult application of long-term culture within a humidified incubator. Some variation in vascular resistance likely originated in the necessity of removing the camera and holder from the pump to change media (Fig. 11A). It is exceedingly unlikely that the camera can be exactly repositioned after the media change such that the fixed fiducial marks were returned to their initial position. The current experimental protocol did not re-establish the initial configuration of the fiducial marks, although any protocol could include a routine where pumping was stopped, the measurement capacitor pressure equilibrated, and the undeformed position of the fixed fiducial marks was re-established. An important feature of the system going forward will be to ensure that the camera holder allows media changes without separating the camera from the pump. Another problem centered on the appearance of bubbles in the pump. Bubbles can be stationary and in a position that is irrelevant to pump function. However, bubbles in sensitive locations can impede pump function. While it is unlikely that a pump design can be entirely insensitive to the appearance of bubbles, incorporation of physical features that trap bubbles in key locations may make the system more robust for long term use in an incubator.

## Acknowledgements

MF was supported by an MIT MathWorks Fellowship. This work was supported by a grant from the National Cancer Institute (U54-CA261694).

## Conflicts of Interest

The authors declare no conflict of interest.

## Figures

**Figure S1.**
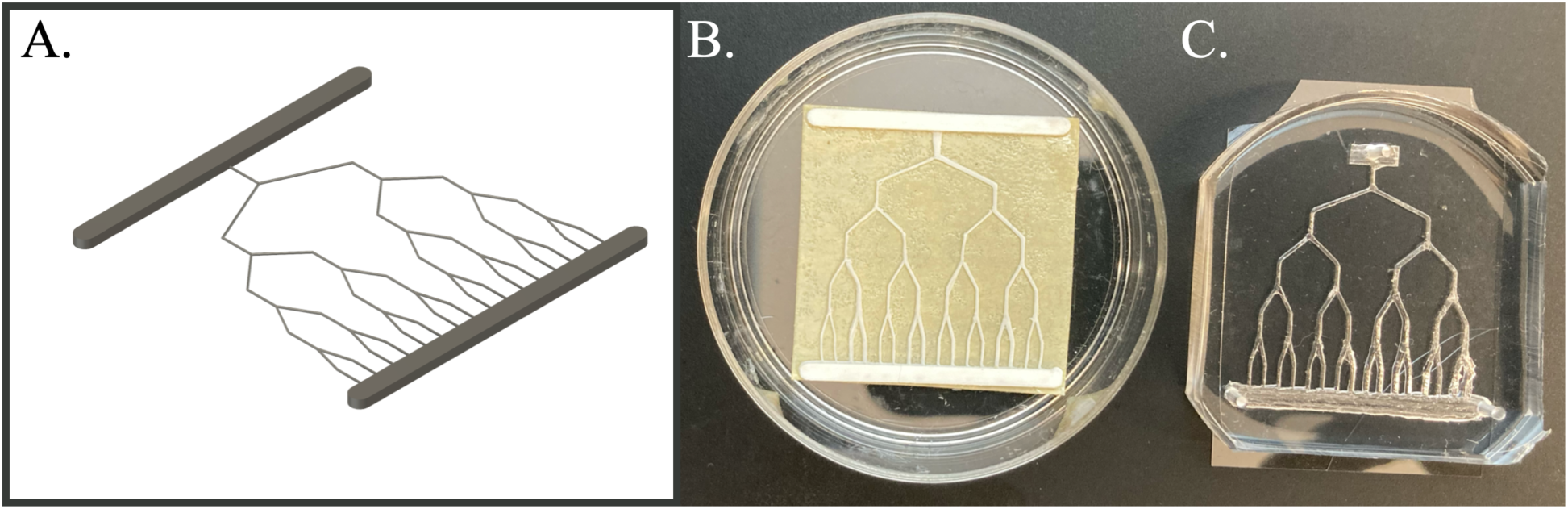
A bifurcating network to model the flow resistance of an interconnected, bifurcating vascular network. (A) 3D printed model showing large reservoirs connected by a network of bifurcating segments. (B) 3D printed network fixed to a petri dish by double-sided tape to for a mold. (C) Model vascular network formed in PDMS and bonded to a coverglass.

**Figure S2.**
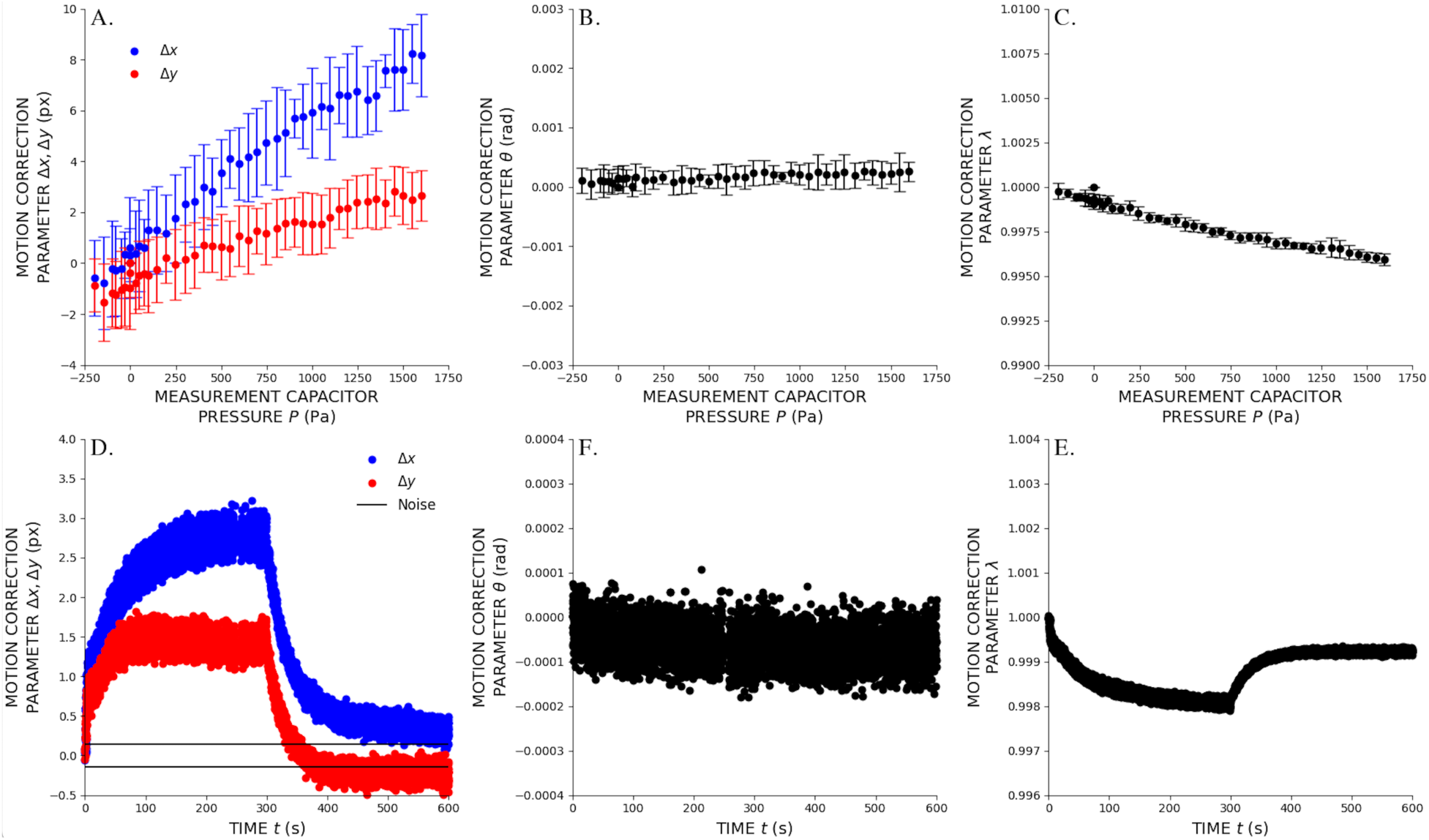
Motion correction parameters computed for each frame captured during calibration of five pumps (*n* = 5). (A) Motion correction parameters Δ*x* (blue symbols) and Δ*y* (red symbols) representing spurious rigid translation of the fiducial marks. (B) Motion correction parameter λ representing uniform dilation. (C) Motion correction parameter θ representing rigid rotation. (D) Motion correction parameters Δ*x* (blue symbols) and Δ*y* (red symbols) computed during loading and unloading of a pump for a representative run driven by an external pressure *P_EX_* = 8.2 kPa. The level of noise is also indicated (black lines). (E) Motion correction parameter λ representing uniform dilation. (F) Motion correction parameter θ representing rigid rotation.

**Figure S3.**
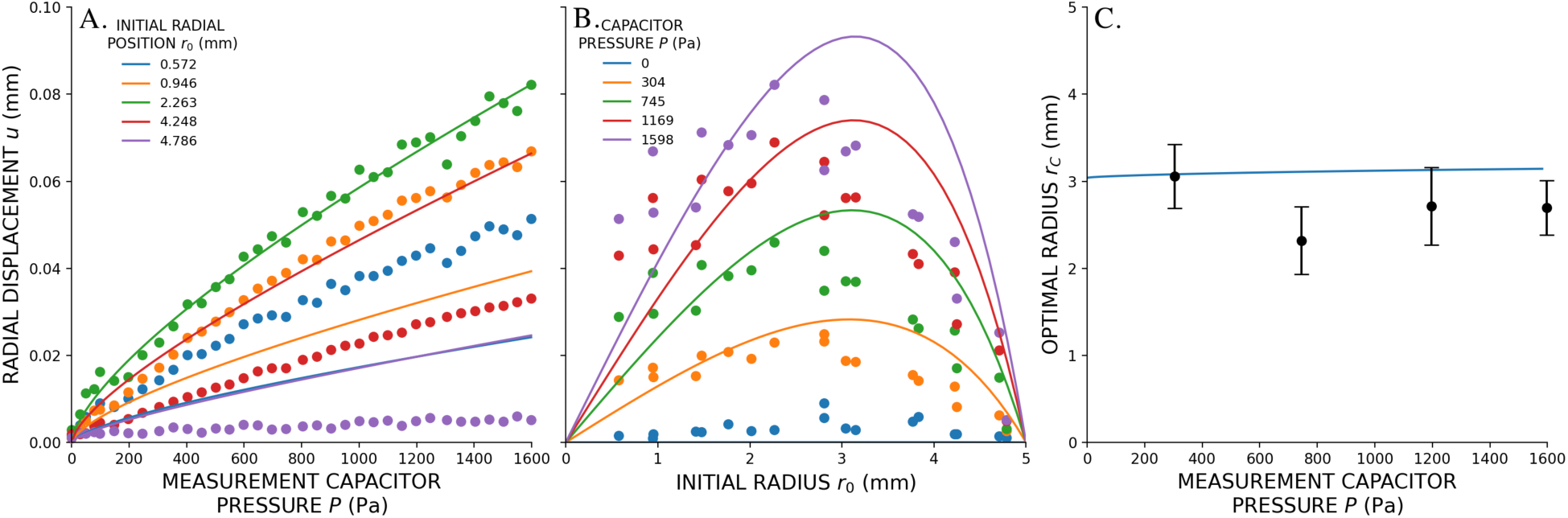
Comparison of experimental data (symbols) to model predictions (lines). (A) The experimental data collected using the optical system (from a single pump subjected to five independent experiments from Figure 5B, with error bars removed for clarity) captured major qualitative features of the radial displacement *u* vs capacitor pressure *P* relationship. The model predicted an increase radial displacement *u* with increased with capacitor pressure *P* and the slight decrease in slope of the *u* vs. *P* relationship with increasing *P*. With *E* = 0.4 MPa, the data and model were in excellent quantitatively agreement for *u* at an initial radial position *r*_0_ = 2.263 mm. Note that the *r*_0_ = 0.572 and 4.786 mm curves are nearly identical. (B) The model predicted the dependence of *u* on both *r*_0_ and *P*. Data from Figure 5C with error bars removed for clarity. (C) The initial radius that maximizes radial displacement *r_C_* for various measurement capacitor pressures. The data include the single pump displayed in (B) and five additional pumps subjected to five independent measurements. The optimal radius is quantitatively close to the model prediction. The optimal radius *r_C_* is very weakly dependent on *P*.

## Supplementary Material

### Spurious Motion

Fiducial marker motion was captured by a camera positioned directly above the capacitor (Fig. 1B). The camera sat on posts of a camera holder that was designed and fabricated in-house. The camera holder was press fit into the upper pump slab. In the absence of membrane deformation, rigid motion of the camera relative to the capacitor would manifest as dilation, rotation, and translation of the fixed fiducial marks. The fixed fiducial marks were used to correct for such motion. It was assumed that relative motion of the camera could result in a uniform dilation characterized by a stretch ratio *λ*, rigid body rotation characterized by an angle *θ*, and rigid translations along the *x*- and *y*-coordinates by displacements Δ*x* and Δ*y*. A fiducial mark initially at position (*X*, *Y*) would be carried by sequential dilation, translation, and rotation to the new position (*x*, *y*) given by

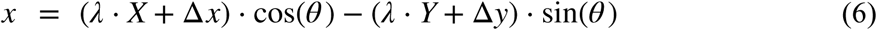

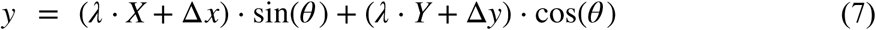

To estimate λ, θ, Δ*x*, and Δ*y* for captured images, the positions of the fixed fiducial marks (*X*_1_, *Y*_1_), (*X*_2_, *Y*_2_), …, (*X_N_f__*, *Y_N_f__*) were determined in a reference image captured with no external pressure. In most cases, all fixed fiducial marks were tracked giving *N_f_* = 6. If one or more fixed fiducial marks were not identified or tracked, *N_f_* < 6. The same fixed fiducial marks were tracked in a subsequent image during pumping (*x*_1_, *y*_1_), (*x*_2_, *y*_2_), …, (*x_N_f__*, *y_N_f__*). If spurious motion was present, *λ*, *θ*, Δ*x*, and Δ*y* were estimated by minimizing the square distance *E*^2^ between the *N_f_* fixed fiducial marks observed during pumping (*x_i_*, *y_i_*) and in the undeformed configuration (*X_i_*, *Y_i_*)

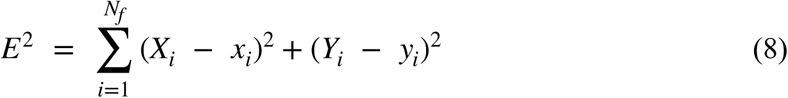

Substituting (6) and (7) into (8) and requiring the partial derivatives taken with respect to *λ*, *θ*, Δ*x*, and Δ*y* all vanish gives the criteria to minimize *E*^2^

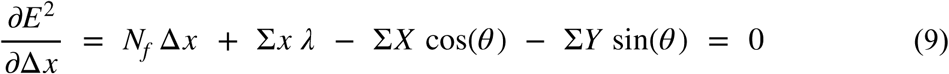

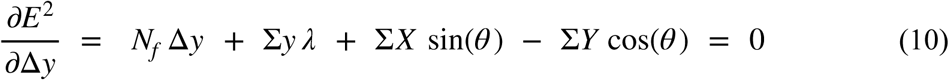

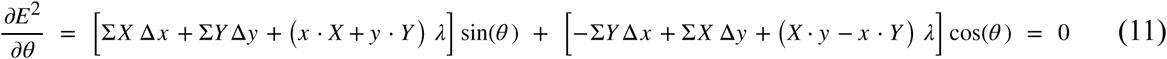

and

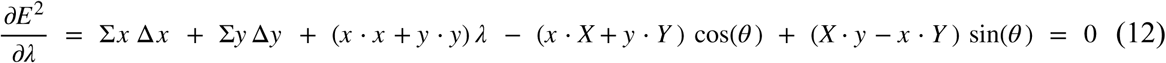

where the notation in Eq. (9) - (12) was simplified using the following substitutions:

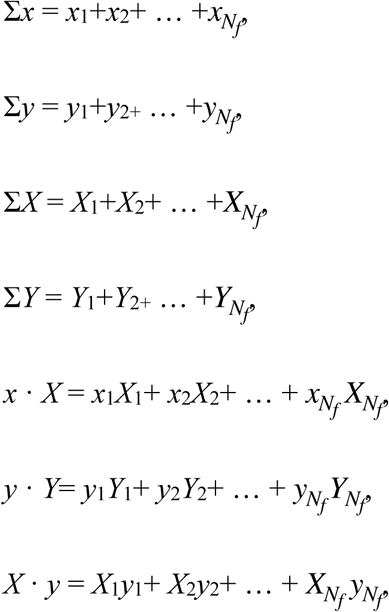

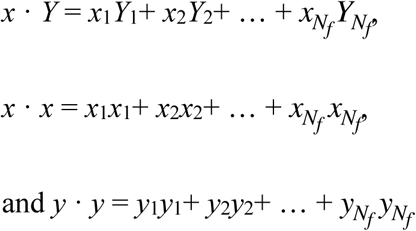

and 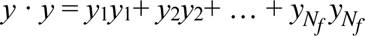

The solution of these four simultaneous equations give expressions for the parameters λ, θ, and Δ*x* and Δ*y* in terms of the positions of the fiducial marks

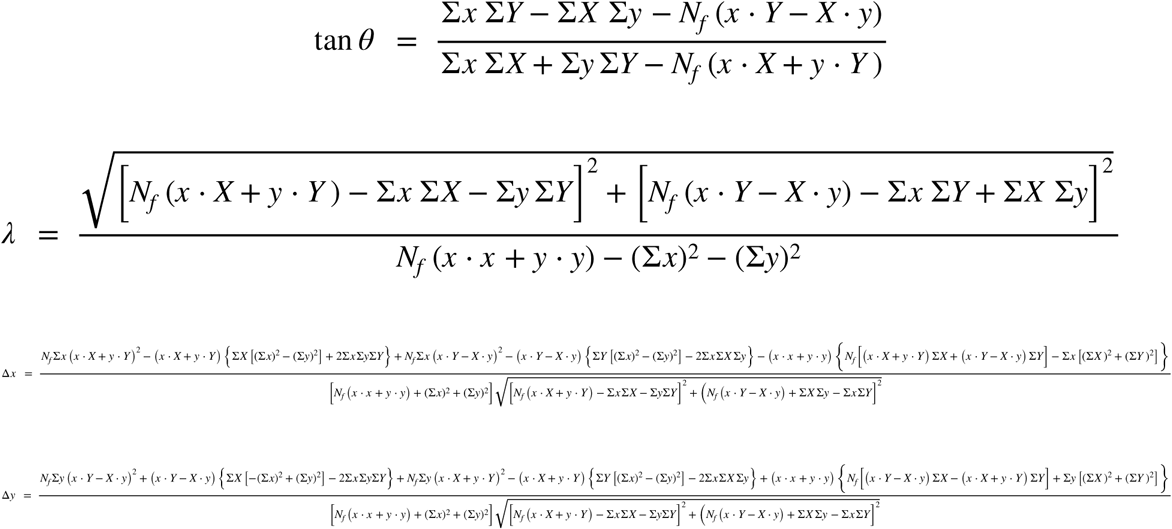

The motion induced by *λ*, *θ*, and Δ*x* and Δ*y* was subtracted from all fiducial marks to give the displacement of fiducial marks attributed to membrane deformation. The corrected positions of the free fiducial marks were used in all subsequent analyses.

The correction parameters Δ*x* and Δ*y* were above the level of system noise suggesting that the fixed fiducial marks exhibited rigid motion relative to their position in the image captured in the initial, unloaded configuration (Fig. S2A). Both Δ*x* and Δ*y* tracked the capacitor pressure (Fig. 8A). This coupling could result from distortion of the pump during pressurization since the pump is made from deformable PDMS. The parameters λ, θ were both very small. This suggest that there was little dilation or rotation of the fixed fiducial marks relative to the initial configuration (Fig. S2B and C).

### Deformation of a thin, elastic membrane subjected to uniform pressure

The mechanical response of a thin elastic membrane subjected to a uniform pressure has been analyzed in detail [cf. 27]. The constitutive behavior of the membrane was assumed to be linearly elastic while the deformation was assumed to be finite. Curiously, the strain-displacement equation for radial strain in the full formulation for finite deformation includes two nonlinear terms but typically only one is retained. Thus, the problem of determining the deformed membrane configuration was nonlinear and required a numerical approach.

Following the approach of Fichter [27], the equilibrium, stress-strain, and strain-displacement equations were reduced to a nonlinear ordinary differential equation in the radial component of the stress resultant and the vertical displacement of the membrane. Radial stress resultant and lateral displacement are expanded in a power series of the radial coordinate of the membrane. The coefficients of each expansion are determined by the boundary condition that displacement at the membrane edge is zero and the requirement that the largest powers of the radial component must satisfy the governing equations. This approach gives good approximations for the deformed membrane shape with a modest number of terms.

Radial displacement *u* increased systematically with capacitor pressure *P* (Fig. S3A). The increase in *u* with increasing *P* was nearly linear, but the slope of the *u*(*P*) curve decreased slightly with increasing *P*. The dependence of slope on *P* was more pronounced at low *u*, so the *u*(*P*) curves showed slight splay. Measurements of radial displacement at various capacitor pressures were in excellent qualitative agreement with the model (Fig. 6B): *u* increased nearly linearly with *P* with evidence of a slight decreasing slope with increasing *P*. Using the known membrane thickness of 0.3 mm and values consistent with the manufacturer’s values for Poisson’s ratio of 0.48 and elastic modulus of 50,000 Pa, the experimental observations and model predictions were in good quantitative agreement. The largest discrepancy between the experimental observations and model predictions occurred for fiducial marks near the capacitor edge. The measured radial displacement was higher than the model predictions. This discrepancy is likely due to misalignment between the fiducial marks and the capacitor.

The radial displacement *u* was zero at the center of the membrane, increased with radial position *r* to a maximum, and decreased to zero at the capacitor edge (*r* = 5 mm) (Fig. S3B). The experimental observations were in remarkably good qualitative and quantitative agreement with the model. The radius that provides the maximum radial displacement for a given pressure was a weak function of Poisson’s ratio and hydrostatic pressure giving values around 0.62 *R* (Fig. S3C).

The model of membrane deformation under uniform pressure is built on two reasonable assumptions: the membrane is linear elastic and the deformation can be large. The model predictions are in excellent agreement with experimental observations suggesting that the model could be a valuable tool for analysis of pumps with imperfect alignment of pump slabs or fiducial marks.

## Notes

### Competing Interest Statement

The authors have declared no competing interest.

## References

1. Chen, M.B., Whisler, J.A., Jeon, J.S., and Kamm, R.D., 2013, “Mechanisms of Tumor Cell Extravasation in an in Vitro Microvascular Network Platform,” Integr Biol (Camb), 5(10), pp. 1262–1271.

2. Moya, M.L., Hsu, Y.H., Lee, A.P., Hughes, C.C., and George, S.C., 2013, “In Vitro Perfused Human Capillary Networks,” Tissue Eng Part C Methods, 19(9), pp. 730–737.

3. Kim, S., Lee, H., Chun, M., and Jean, N. L., 2013, “Engineering of Functional, Perfusable 3D Microvascular Networks on a Chip,” Lab on a Chip, 13(8), pp. 1489–1500.

4. Myers, D. R. And Lam, W. A., 2021, “Vascularized Microfluidics and Their Untapped Potential for Discovery in Diseases of the Microvasculature,” Ann Rev Biomed Eng, 23, pp. 407–432.

5. Bersini, S. and Moretti, M., 2015, “3D Functional and Perfusable Microvascular Networks for Organotypic Microfluidic Models,” J Mater Sci Mater Med, 26(5), pp. 180.

6. Dessalles, C.A., Leclech, C., Castagnino, A., and Barakat, A.I., 2021, “Integration of Substrate- and Flow-Derived Stresses in Endothelial Cell Mechanobiology,” Commun Biol, 4(1), pp. 764.

7. Gifre-Renom, L., Daems, M., Luttun, A., and Jones, E.A.V., 2022, “Organ-Specific Endothelial Cell Differentiation and Impact of Microenvironmental Cues on Endothelial Heterogeneity,” Int J Mol Sci, 23(3), pp. 1477.

8. Chien, S., 2007, “Mechanotransduction and Endothelial Cell Homeostasis: The Wisdom of the Cell,” Am J Physiol Heart Circ Physiol, 292(3), pp. H1209–1224.

9. Offeddu, G.S., Serrano, J.C., Chen, S.W., Shelton, S.E., Shin, Y., Floryan, M., and Kamm, R.D., 2021, “Microheart: A Microfluidic Pump for Functional Vascular Culture in Microphysiological Systems,” J Biomech, 119, pp. 110330.

10. Mosadegh, B., Kuo, C.H., Tung, Y.C., Torisawa, Y.S., Bersano-Begey, T., Tavana, H., and Takayama, S., 2010, “Integrated Elastomeric Components for Autonomous Regulation of Sequential and Oscillatory Flow Switching in Microfluidic Devices,” Nat Phys, 6(6), pp. 433–437.

11. King, P., The Camera Module Guide, in Take Pictures and Capture Video with your Raspberry Pi, P. King, Editor., Select Publisher Services, Ltd: Cambridge. p. 124.

12. Fabry, B., Maksym, G.N., Shore, S.A., Moore, P.E., Reynold A. Panettieri, J., Butler, J.P., and Fredberg, J.J., 2001, “Time Course and Heterogeniety of Contractile Responses in Cultured Human Airway Smooth Muscle Cells,” J Appl Physiol, 91(2), pp. 986–994.

13. Shin, Y., Han, S., Jeon, J.S., Yamamoto, K., Zervantonakis, I.K., Sudo, R., Kamm, R.D., and Chung, S., 2012, “Microfluidic Assay for Simultaneous Culture of Multiple Cell Types on Surfaces or within Hydrogels,” Nat Protoc, 7(7), pp. 1247–1259.

14. Blazeski, A., Floryan, M.A., Fajardo-Ramirez, O.R., Meibalan, E., Ortiz-Urbina, J., Angelidakis, E., Shelton, S.E., Kamm, R.D., and Garcia-Cardena, G., 2024, “Engineering Microvascular Networks Using a KLF2 Reporter to Probe Flow-Dependent Endothelial Cell Function,” Biomaterials, 311, pp. 122686.

15. Haase, K., Piatti, F., Marcano, M., Shin, Y., Visone, R., Redaelli, A., Rasponi, M., and Kamm, R.D., 2022, “Physiologic Flow-Conditioning Limits Vascular Dysfunction in Engineered Human Capillaries,” Biomaterials, 280, pp. 121248.

16. Cherubini, M., Erickson, S., Padmanaban, P., Haberkant, P., Stein, F., Beltran-Sastre, V., and Haase, K., 2023, “Flow in Fetoplacental-Like Microvessels in Vitro Enhances Perfusion, Barrier Function, and Matrix Stability,” Sci Adv, 9(51), pp. eadj8540.

17. Moccia, C., Cherubini, M., Fortea, M., Akinbote, A., Padmanaban, P., Beltran-Sastre, V., and Haase, K., 2023, “Mammary Microvessels Are Sensitive to Menstrual Cycle Sex Hormones,” Adv Sci (Weinh), 10(35), pp. e2302561.

18. Floryan, M., Cambria, E., Blazeski, A., Coughlin, M.F., Wan, Z., Offeddu, G.S., Vinayak, V., Kant, A., Shenoy, V., and Kamm, R.D., 2025, “Remodeling of Self-Assembled Microvascular Networks Under Long Term Flow,” npj J Biol Phys and Mech, (In Press)

19. Hajal, C., Ibrahim, L., Serrano, J.C., Offeddu, G.S., and Kamm, R.D., 2021, “The Effects of Luminal and Trans-Endothelial Fluid Flows on the Extravasation and Tissue Invasion of Tumor Cells in a 3D in Vitro Microvascular Platform,” Biomaterials, 265, pp. 120470.

20. Wan, Z., Zhang, S., Zhong, A.X., Xu, L., Coughlin, M.F., Pavlou, G., Shelton, S.E., Nguyen, H.T., Hirose, S., Kim, S., Floryan, M.A., Barbie, D.A., Hodi, F.S., and Kamm, R.D., 2024, “Transmural Flow Upregulates PD-L1 Expression in Microvascular Networks,” Adv Sci, 11(26), pp. e2400921.

21. Shirure, V.S., Lezia, A., Tao, A., Alonzo, L.F., and George, S.C., 2017, “Low Levels of Physiological Interstitial Flow Eliminate Morphogen Gradients and Guide Angiogenesis,” Angiogenesis, 20(4), pp. 493–504.

22. Byun, C.K., Abi-Samra, K., Cho, Y.K., and Takayama, S., 2014, “Pumps for Microfluidic Cell Culture,” Electrophoresis, 35(2-3), pp. 245–257.

23. Kaarj, K. and Yoon, J.Y., 2019, “Methods of Delivering Mechanical Stimuli to Organ-on-a-Chip,” Micromachines (Basel), 10(10),

24. Schimek, K., Busek, M., Brincker, S., Groth, B., Hoffmann, S., Lauster, R., Lindner, G., Lorenz, A., Menzel, U., Sonntag, F., Walles, H., Marx, U., and Horland, R., 2013, “Integrating Biological Vasculature into a Multi-Organ-Chip Microsystem,” Lab on a Chip, 13(18), pp. 3588–3598.

25. Uesugi, K., Nishiyama, K., Hirai, K., Inoue, H., Sakurai, Y., Yamada, Y., Taniguchi, T., and Morishima, K., 2020, “Survival Rate of Cells Sent by a Low Mechanical Load Tube Pump: The “Ring Pump”,” Micromachines (Basil), 11(4), pp. 447.

26. Pozdin, V.A., Erb, P.D., Downey, M., Rivera, K.R., Daniele, M., 2021, “Monitoring of Random Microvessel Network Formation by In-Line Sensing of Flow Rates: A Numerical and In Vitro Investigation,” Sensors and Actuators A: Physical, 331, pp. 112970.

27. Fichter, W.B., 1997, “Some Solutions for the Large Deflections of Uniformly Loaded Circular Membranes,” NASA Technical Paper 3658, July, pp. 1-20.

